# Humans adaptively select different computational strategies in different learning environments

**DOI:** 10.1101/2023.01.27.525944

**Authors:** Pieter Verbeke, Tom Verguts

**Affiliations:** Department of Experimental Psychology; Ghent University; Ghent; B9000

**Keywords:** Adaptive model selection, Hierarchical learning, Reinforcement learning, Cognitive flexibility

## Abstract

The Rescorla-Wagner rule remains the most popular tool to describe human behavior in reinforcement learning tasks. Nevertheless, it cannot fit human learning in complex environments. Previous work proposed several hierarchical extensions of this learning rule. However, it remains unclear when a flat (non-hierarchical) versus a hierarchical strategy is adaptive, or when it is implemented by humans. To address this question, current work applies a nested modelling approach to evaluate multiple models in multiple reinforcement learning environments both computationally (which approach performs best) and empirically (which approach fits human data best). We consider ten empirical datasets (*N* = 407) divided over three reinforcement learning environments. Our results demonstrate that different environments are best solved with different learning strategies; and that humans adaptively select the learning strategy that allows best performance. Specifically, while flat learning fitted best in less complex stable learning environments, humans employed more hierarchically complex models in more complex environments.

## Introduction

Humans and other primates are remarkably flexible in adapting to changing environments. For this purpose, they should keep track of contingencies between stimuli, actions, and rewards. To date, such reward contingency learning is often described by what are called flat (i.e., non-hierarchical) learning algorithms. Perhaps the most popular instance of flat learning is the Rescorla-Wagner (RW) model (Rescorla & Wagner, 1972; Sutton & Barto, 1998). This RW learning rule allows to describe the behavioral learning curve (increase in accuracy over time) by estimating a single learning rate parameter. Importantly, flat learning algorithms learn only one set of stimulus-action mappings (further referred to as rule set). Hence, each stimulus is associated to exactly one optimal action. This is limited as a model of human cognition since the most optimal action given a certain stimulus often depends on the current goal or context (i.e., the task). For example, while one needs to drive on the left side of the road in the UK, the European mainland requires driving on the right side of the road. A flat learning agent would need to constantly overwrite/ relearn this information when it switches contexts (countries).

Thus, although flat learning works well in stable and simple environments it has often proven to be insufficient to describe adaptive human behavior in more volatile or complex environments (Bai et al., 2014; McGuire et al., 2014; Verbeke, Ergo, De Loof, & Verguts, 2021; Wilson et al., 2013). An empirical example are probabilistic reversal learning tasks (Cools et al., 2002; Izquierdo et al., 2017; Verbeke, Ergo, De Loof, & Verguts, 2021). In these tasks, an agent faces two contradictory goals. On the one hand, the agent should be flexible to deal with sudden reversals of the stimulus-action-reward contingencies (plasticity). On the other hand, the agent should also be robust to the noise that is induced by the probabilistic feedback (stability). In contrast to humans (Nassar & Troiani, 2021), flat learning algorithms cannot achieve a balance between stability and plasticity in a dynamic manner (see also Grossberg, 1987). This is because high learning rates would lead the agent to “chase the noise” and adapt behavior when the contingencies did not change but low learning rates would significantly decrease the flexibility of the agent in adapting to changing contingencies. As a result, several hierarchical extensions of the RW rule have been proposed to better describe human flexibility (Behrens et al., 2007; Bouchacourt et al., 2022; Mathys et al., 2011; McGuire et al., 2014; Verbeke, Ergo, De Loof, & Verguts, 2021). Across several reinforcement learning environments, the current paper evaluates when flat versus hierarchical learning is computationally beneficial for performance, when it fits human data, and how these two factors relate to one another.

### Hierarchical extensions to the Flat model

Current work aims to investigate when hierarchically extended models are beneficial. For this purpose, a nested modelling approach is applied (Grainger & Jacobs, 1996; Perry et al., 2007; Vidal & Durfee, 1998). Specifically, three hierarchical extensions to the RW model are considered. Each extension is implemented by adding a single free parameter to the model. In this sense, each extension is equally complex in computational terms. Our nested modelling approach allows to evaluate the added value of each single extension but also how the extensions interact.

A first popular extension to the flat learning approach is to introduce a hierarchically higher parameter which allows to adaptively change lower-level parameters, such as the learning rate (Bai et al., 2014; Behrens et al., 2007; McGuire et al., 2014). For instance, in reversal learning, it can be beneficial to increase the learning rate if uncertainty in the environment (measured e.g., by prediction errors) is high (to increase flexibility); and decrease the learning rate if uncertainty is low to shield learning against noise in stable periods (as in the Kalman filter; Kruschke, 2008). Adaptive learning rates can be implemented in multiple ways, including adding one parameter to the classic reinforcement learning rule (Bai et al., 2014); using Bayesian learning frameworks (Mathys et al., 2011); using a mixture of learning rules (Wilson et al., 2013); making learning rate dependent on prediction error (Silvetti et al., 2017); and setting a different learning rate for each individual synapse (Schweighofer & Arbib, 1998). Current work aimed to limit the number of additional parameters and used an approach that could easily be integrated with the other hierarchical extensions. Therefore, a Hybrid parameter (see Bai et al., 2014) was implemented which increases the learning rate in response to increasing uncertainty in the environment.

A second extension consists of storing multiple rule sets and adaptively switch between these sets when there is a large change in stimulus-action reward contingencies (Bouchacourt et al., 2022; Razmi & Nassar, 2022; Verbeke, Ergo, De Loof, & Verguts, 2021; Verbeke & Verguts, 2019). This extension implements a hierarchical architecture for learning. At the lower level, multiple rule sets are learned. At the higher level, the agent decides when to switch between these rule sets. Thus, if there are multiple sufficiently distinctive sets of contingencies that are frequently revisited over the time course of the task (as in reversal learning tasks), a hierarchical agent can learn multiple (robust) rule sets and then on a hierarchically higher level decide when to switch between rule sets. Current work implements the switch decision as an accumulation-to-bound process in which negative feedback (i.e., absence of reward) is accumulated over time. When a bound is reached, the agent switches to another rule set. For this purpose, a Cumulation parameter is implemented which specifies the speed at which negative feedback is accumulated for the switch decision (Verbeke, Ergo, De Loof, & Verguts, 2021; Verbeke & Verguts, 2019).

Often, although not always, models with multiple rule sets also implement learning at the hierarchically higher level (Botvinick et al., 2009; Collins & Frank, 2013; Eckstein & Collins, 2020; Verbeke & Verguts, 2019). This allows to switch between different rule sets more adaptively. For instance, one could learn to map task contexts to rule sets. As a result, one could infer from context which rule set to switch to (Botvinick et al., 2009; Collins & Frank, 2013; Eckstein & Collins, 2020). Additionally, learning on the hierarchical level could allow adapting to changing levels of noise in reward feedback and hence learning when to switch (Foucault & Meyniel, 2021; Verbeke & Verguts, 2019; L. Q. Yu et al., 2021). Specifically, while with low levels of feedback noise, an agent might decide to switch to another rule set every time it receives negative feedback, high levels of feedback noise would require agents to be more conservative about switching since negative feedback might be caused by noise instead of an actual change in stimulus-action reward contingencies (A. J. Yu & Dayan, 2005). Thus, the weight of negative feedback in the decision to switch might be decreased. Current work implements learning on the hierarchically higher level to adapt the weight of negative feedback in the accumulation-to-bound process that determines when to switch (Verbeke, Ergo, De Loof, & Verguts, 2021; Verbeke & Verguts, 2019).

As illustrated in Figure 1A, six nested models are created by combining three possible extensions: adaptive learning rate, multiple rule sets, and hierarchical learning. All models have the Flat model as base but add a different set of hierarchical extensions by allowing a different combination of additional parameters to be different from zero. Notice that hierarchical learning builds on the multiple rule sets extension. Hence, the two combinations with hierarchical learning but no multiple rule sets are impossible, leaving just 2^3^ – 2 = 6 possible models.

**Figure 1.**
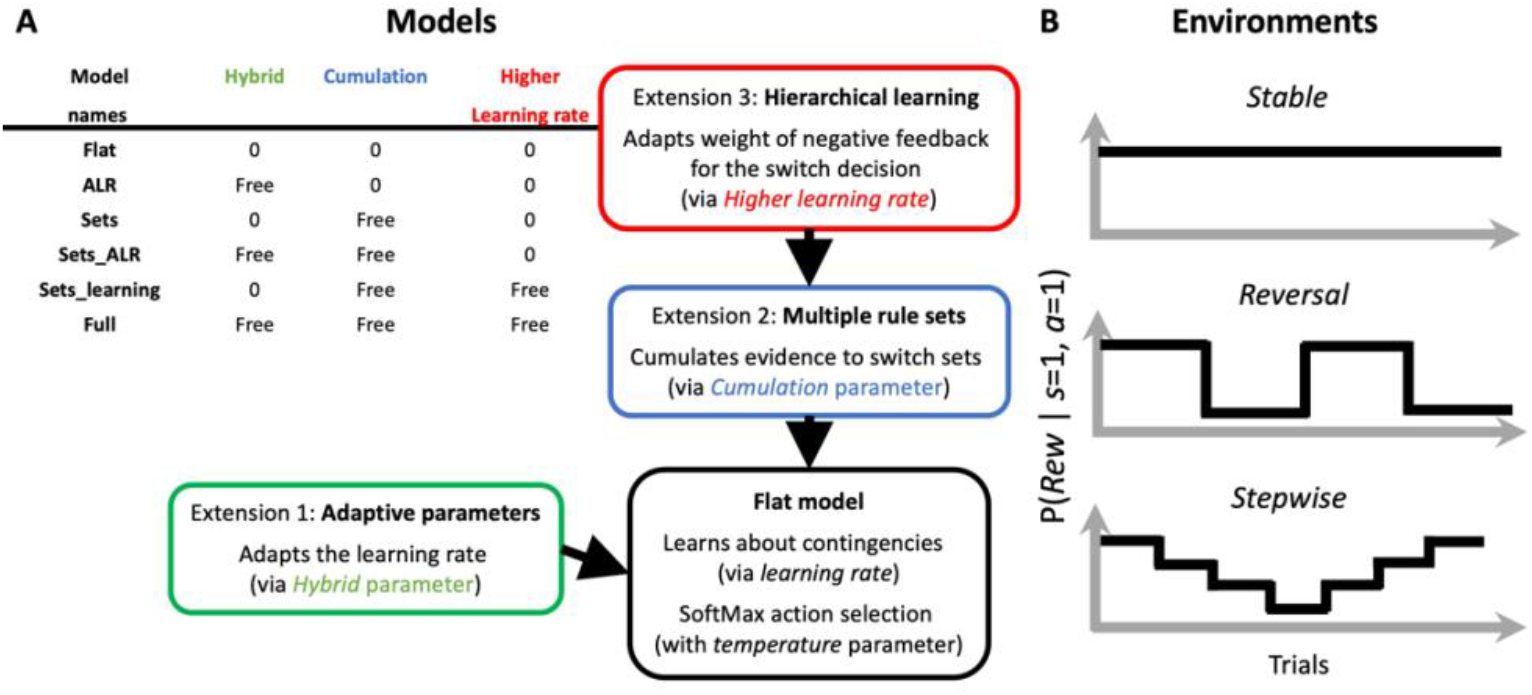
Study set-up. We evaluate multiple (nested) models in multiple environments. *A: The models.* A schematic illustration of the three extensions to the Flat model. Each extension is implemented by adding a single parameter (indicated in red, blue or green). The model list demonstrates how different combinations of these extensions result in six possible models. Here, parameters are freely estimated on the data or fixed to zero. Parameters that are fixed to zero do not influence model behavior. Each model is an extension of the Flat model, so they all have a learning rate and temperature parameter that is freely estimated (see Methods for details). *B: The reinforcement learning environments.* This figure provides a schematic illustration of how stimulus-action-reward contingencies could evolve in each of the three environments.

Another increasingly popular computational approach to describe human behavior are artificial neural networks (Ma & Peters, 2020). Although these networks have many more parameters and are therefore also much more difficult to interpret, they often reach a better fit or performance than classic cognitive models. Hence, they provide a good benchmark to evaluate cognitive models (Ji-An et al., 2023). Moreover, investigating neural network activation in the hidden layer(s) allows to explore how the neural network organizes task representations for performance. Therefore, we also implemented a neural network. More specifically, a Gated Recurrent Unit (GRU; Chung et al) was trained and used as a normative model to provide a benchmark for (classic) cognitive model performance but also provide insights in optimal cognitive strategies for each environment.

### The environments

Traditionally, models are evaluated by measuring the goodness-of-fit on a single behavioral dataset. By comparing the goodness-of fit across two or more models, one can determine which model provides the best explanation for the underlying cognitive processes. Nevertheless, previous work has argued that such findings generalize poorly across tasks (Eckstein et al., 2021, 2022). Whether and which hierarchical extensions are needed, may depend on the environment. As a result, one model might fit better than the alternative in some task, but the alternative model might fit better in another task. Also, it often remains elusive whether and how better behavioral fit relates to better performance. Therefore, current work proposes a novel approach for model evaluation (see also van den Berg et al., 2012). Instead of considering a single dataset and one particular task, we compare models across multiple datasets and reinforcement learning tasks. Furthermore, we evaluate models both computationally, in terms of task performance, as well as empirically in terms of goodness-of-fit. Such an in-depth model comparison allows to investigate how learning strategies differ across tasks and how this relates to behavioral performance.

One popular approach to study reinforcement learning is via two-armed bandit tasks (Moran et al., 2021; Noonan et al., 2010; Silvetti et al., 2011; Sutton & Barto, 1998). Here, on each trial there is one optimal action yielding the highest probability of reward and one suboptimal action yielding a lower probability of reward. Subjects thus need to learn the optimal action for each stimulus. Two-armed bandit tasks can be made more challenging by implementing more dynamic reinforcement environments. Current work evaluates multiple models in three task environments that differ in terms of the reinforcement dynamics. An overview of the reinforcement dynamics in each environment is given in Figure 1B.

In the Stable environment, one rule set needs to be learned, whose contingencies do not change. Hence, the identity of the optimal action (*a_optimal_*) as well as the probability of reward (*Rew*) given this optimal action (P(*Rew* | *a_optimal_*)) remain stable over time (over trials). To evaluate goodness-of-fit, we fitted all models on three publicly available behavioral datasets in the Stable environment (Goris et al., 2021; Huycke et al., 2021; Xia et al., 2021). Although P(*Rew* | *a_optimal_*) was stable within one dataset, it varied between 70% and 100% across datasets. To evaluate model performance, we simulated all cognitive models and the GRU network on all three stable datasets.

In the Reversal environment, two orthogonal rule sets are frequently revisited during the time course of the task. Thus, while P(*Rew* I *a_optimal_*) remains stable over time, the identity of *a_optimal_* regularly reverses during the time course of the task. We found four publicly available two-armed bandit tasks in the Reversal environment (Goris et al., 2021; Liu et al., 2022; Mukherjee et al., 2020; Verbeke & Verguts, 2019). Again, P(*Rew* I *a_optimal_*) was stable within a dataset but varied across datasets. Across the four publicly available datasets, P(*Rew* I *a_optimal_*) varied from 70 to 90%. Notice that the deterministic version (P(*Rew* I *a_optimal_*) = 100%) of the reversal learning task is a special case because there is no feedback noise. As a result, flat learning will experience less problems since a constant high learning rate cannot lead to chasing the noise when noise is absent. There is no stability problem. To control for this special case, we considered it necessary to include such a dataset. Nevertheless, we did not find a dataset that was available and suited for comparison (in terms of design, subjects, …) with the other datasets. For this purpose, an additional (not previously published) dataset was collected in the Reversal environment with P(*Rew* I *a_optimal_*) = 100% (see Methods). Hence, after this additional data collection, also the datasets in the Reversal environment varied from 70 to 100% in terms of P(*Rew* I *a_optimal_*). To evaluate model performance, we again performed simulations on each Reversal dataset for all cognitive models as well as the GRU network.

In the Stepwise environment, the identity of the optimal action also changed over time. However, another layer of complexity was added in these datasets since also P(*Rew* I *a_optimal_*) could vary within the dataset. In the two Stepwise datasets (Cohen et al., 2007; Hein et al., 2021), P(*Rew* I *a_optimal_*) varied from 50 to 90%. Again, models and the GRU network were simulated on both datasets.

In sum, we provide an extensive model comparison over multiple reinforcement learning environments and empirical datasets. This allows to investigate when and which hierarchical extensions to the flat learning RW model provide a benefit in terms of empirical fit and performance. Six nested models are created by combining three possible extensions: adaptive learning rate, multiple rule sets, and hierarchical learning. These models are tested on the Stable, Reversal and Stepwise environments. For this purpose, we investigated ten datasets (total *N* = 407) across the three types of environments. For each environment we aim to extract the simplest (fewest parameters) model that performs best and that fits empirical data best. Hence, the added value of each model extension is evaluated both computationally in terms of model performance, as well as empirically in terms of model fit.

## Materials and methods

### The cognitive models

We evaluate three hierarchical extensions to the Flat RW model: adaptive learning rates, multiple rule sets and hierarchical learning. As demonstrated in Fig. 1A, six possible (nested) models can be constructed by making different combinations of these extensions.

All models had the Flat model as base. Here, stimulus-action-reward contingencies are learned via the RW learning rule. Thus, the expected reward of one stimulus-action pair (*Q*(*s*,*a*)), is updated on each trial (*t*) by

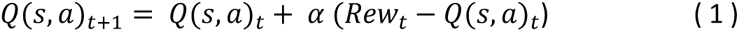

in which α is the learning rate and *Rew_t_* represents the reward received on trial *t*. Note that only the *Q* associated with the presented stimulus and given action is updated on every trial.

For action selection, a SoftMax decision rule is used,

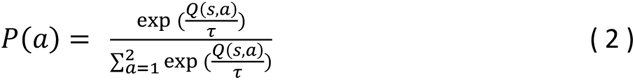

Here, τ is a temperature parameter which controls the degree of exploration. With higher values of τ, there is a higher probability of selecting an action that does not have the highest expected reward.

One possible hierarchical extension of the flat learning approach is to implement an adaptive lower-level parameter. Here, we adopt the approach of Bai et al. (2014) to adapt the learning rate. Specifically,

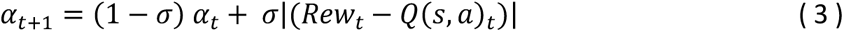

in which we call σ the hybrid parameter. This hybrid parameter allows to continually update the learning rate parameter in both positive and negative directions. Specifically, Equation (3) implies that the learning rate increases on trials with strong prediction errors (*Rew_t_* – *Q*(*s, a*)*_t_*) and decreases with a factor σ otherwise.

The second extension to flat learning considers multiple rule sets (*rs*). Therefore, the expected reward of stimulus-action pairs are learned for each rule set separately (*Q*(*s, a, rs*) instead of *Q*(*s,a*)). Which rule set initially has control of behavior is determined at random on the first trial. In line with previous work (Verbeke, Ergo, De Loof, & Verguts, 2021; Verbeke & Verguts, 2019), an accumulation-to-bound process determines when the model switches to the other rule set. For this purpose, a Cumulation parameter (γ) is implemented which specifies the speed at which negative feedback is accumulated for the switch decision (Verbeke, Ergo, De Loof, & Verguts, 2021; Verbeke & Verguts, 2019). Specifically, Switch evidence *S_t_* accumulates via

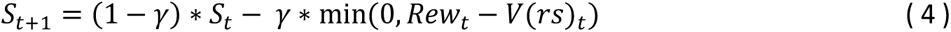

in which *V*(*rs*)_t_ is the expected reward associated with rule set *rs* chosen at trial *t*. Since current work considers two-arm bandit tasks with two rule sets, we initialized the expected reward at .5. On every trial when the reward is smaller than *V*(*rs*)_t_ (that is, on unrewarded trials), the accumulated switch evidence in *S* increases and decreases otherwise. Here, the reward was either 1 (reward) or 0 (no reward) and the γ parameter had bounds [0, 1]. If there is no hierarchical learning, *V*(*rs*) does not change and hence is fixed at .5. Thus, in absence of hierarchical learning, the asymptote of Equation (4) equals .5. Therefore, we implemented that the agent switches to another rule set if the switch evidence *S_t_* reaches a threshold value just below that value (.49).

Note that the model can adaptively switch with a fixed *V(rs*). Nevertheless, in some situations, adding a learning rule for *V*(*rs*) can be beneficial. For instance, when feedback noise decreases (as sometimes happens in Stepwise environments), it might be adaptive to increase the weight of negative feedback (*V*(*rs*)) since it is more likely that this negative feedback was the consequence of a change in the contingencies than of noise in the environment. Hence, we can add RW learning at the hierarchically higher level as well via

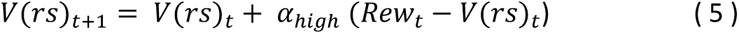

where α*_high_* is the hierarchically higher learning rate (see also Verbeke, Ergo, De Loof, & Verguts, 2021; Verbeke & Verguts, 2019). As illustrated in Fig. 1A, six possible models can be constructed based on the three extensions.

### The GRU network

Additionally, a GRU network was implemented to function as a benchmark for the six cognitive models. At each trial, this network received three inputs: the stimulus of the previous trial (*s_t-1_*), the rewarded response of the previous trial (*r_rew_t-1_*) and the stimulus of the current trial (*s_t_*). Each input was delivered as a one-hot encoded vector. These inputs were provided to a GRU layer. The GRU layer had 5 nodes (*h*). As is implemented by default in Keras (Chollet et al., 2015), these nodes were updated by using a softmax activation function for the reset and update gates and a tanh function for the new gates. Recurrent weights were initialized orthogonally and a glorot uniform kernel was used to initialize feedforward weights. The output layer consisted of two output neurons with a softmax activation function.

### The environments

As described in the Introduction, the current work considers three reinforcement learning environments. All environments can be described as two-armed bandit tasks but differed in the nature of the reinforcement dynamics (See Fig. 1B). We collected ten existing datasets (total *N* = 407) across three types of environments via osf.io, or by contacting the corresponding authors. This resulted in three datasets in the Stable environment (Goris et al., 2021; Huycke et al., 2021; Xia et al., 2021), four in the Reversal environment (Goris et al., 2021; Liu et al., 2022; Mukherjee et al., 2020; Verbeke, Ergo, De Loof, & Verguts, 2021) and two in the Stepwise environment (Cohen et al., 2007; Hein et al., 2021). We collected one additional dataset in the Reversal environment via prolific (www.prolific.co) for reasons that we described in the Introduction. Note that the probability to obtain reward given an optimal action (P(*Rew* I *a_optimal_*)) differed across datasets and simulations. In the datasets we consider, it is also the case that P(*Rew* I *a_suboptimal_*) = 1 – P(*Rew* I *a_optimal_*). In Table 1, we present a summary of all datasets. A more detailed description of each dataset and environment is provided below.

**Table 1.**
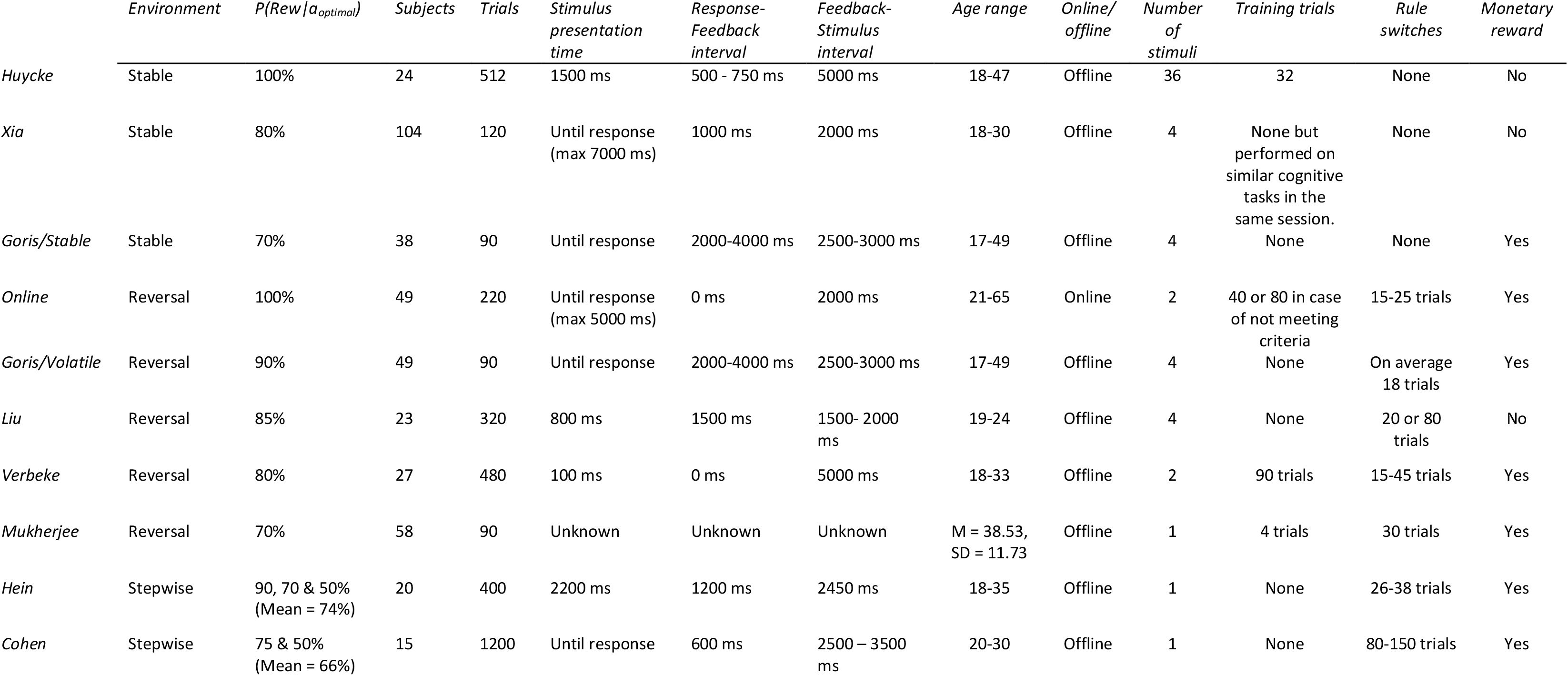
Detailed information on datasets.

|  | *Environment* | *P(Rew\|$a_{optimal}$)* | *Subjects* | *Trials* | *Stimulus presentation time* | *Response-Feedback interval* | *Feedback-Stimulus interval* | *Age range* | *Online/offline* | *Number of stimuli* | *Training trials* | *Rule switches* | *Monetary reward* |
| --- | --- | --- | --- | --- | --- | --- | --- | --- | --- | --- | --- | --- | --- |
| *Huycke* | Stable | 100% | 24 | 512 | 1500 ms | 500 - 750 ms | 5000 ms | 18-47 | Offline | 36 | 32 | None | No |
| *Xia* | Stable | 80% | 104 | 120 | Until response (max 7000 ms) | 1000 ms | 2000 ms | 18-30 | Offline | 4 | None but performed on similar cognitive tasks in the same session. | None | No |
| *Goris/Stable* | Stable | 70% | 38 | 90 | Until response | 2000-4000 ms | 2500-3000 ms | 17-49 | Offline | 4 | None | None | Yes |
| *Online* | Reversal | 100% | 49 | 220 | Until response (max 5000 ms) | 0 ms | 2000 ms | 21-65 | Online | 2 | 40 or 80 in case of not meeting criteria | 15-25 trials | Yes |
| *Goris/Volatile* | Reversal | 90% | 49 | 90 | Until response | 2000-4000 ms | 2500-3000 ms | 17-49 | Offline | 4 | None | On average 18 trials | Yes |
| *Liu* | Reversal | 85% | 23 | 320 | 800 ms | 1500 ms | 1500- 2000 ms | 19-24 | Offline | 4 | None | 20 or 80 trials | No |
| *Verbeke* | Reversal | 80% | 27 | 480 | 100 ms | 0 ms | 5000 ms | 18-33 | Offline | 2 | 90 trials | 15-45 trials | Yes |
| *Mukherjee* | Reversal | 70% | 58 | 90 | Unknown | Unknown | Unknown | M = 38.53, SD = 11.73 | Offline | 1 | 4 trials | 30 trials | Yes |
| *Hein* | Stepwise | 90, 70 & 50% (Mean = 74%) | 20 | 400 | 2200 ms | 1200 ms | 2450 ms | 18-35 | Offline | 1 | None | 26-38 trials | Yes |
| *Cohen* | Stepwise | 75 & 50% (Mean = 66%) | 15 | 1200 | Until response | 600 ms | 2500 – 3500 ms | 20-30 | Offline | 1 | None | 80-150 trials | Yes |

### The datasets

#### Huycke et al., 2021 (Stable 100% reward)

In this dataset (Huycke et al., 2021), subjects had to learn mappings between abstract stimuli and left or right responses. These mappings remained stable across the entire task. Once the subject gave a response, feedback was presented (correct / incorrect). Here, P(*Rew* I *a_optimal_*) = 100%. In total, there were 36 unique stimuli which were presented in groups of 4 stimuli per block. Thirty-two stimuli were only repeated 8 times (in one block); the other 4 stimuli were repeated 64 times (8 times across 8 blocks). Subjects did not receive monetary reward based on performance. All six models were fitted for all subjects (*N* = 24) across all trials (*Trials* = 512). *Xia et al., 2021 (Stable 80% reward)*.

In this dataset (Xia et al., 2021), subjects had to learn which butterfly preferred which flower. These preferences remained stable across the entire task. Two flowers were repeated every trial. The locations of the two flowers (left or right) were randomized over trials. On each trial, one of four possible butterflies was presented, and the subject had to direct the butterfly to the left or right flower with a video game controller. Each butterfly was repeated 30 times across 4 blocks. After a response, feedback was presented (win / lose). Here, P(*Rew* I *a_optimal_*) = 80%. Subjects did not receive monetary reward based on performance. The original study was interested in age-related differences in reinforcement learning. Therefore, there was a strong variability in age. For comparison with other datasets, we fitted the models on the group of subjects that reached adulthood (age >= 18 years). Additionally, we excluded 3 subjects who scored below chance level. As a result, all six models were fitted on *N* = 104 subjects across all trials (*Trials* = 120).

#### Goris et al., 2021 (Stable 70% reward)

In this dataset (Goris et al., 2021), subjects were presented on each trial with a set of 2 pictures and had to use a left or right button press to select respectively the left or right picture. Two sets of pictures were randomly presented across trials. After each response, feedback was presented in the form of a picture of money that was crossed when feedback was negative. Subjects received a small monetary reward based on performance. The full task consisted of three different conditions. Here, we selected what was called the stationary low noise condition. To avoid carry-over from previous conditions, we only analyzed data of subjects who started with our condition of interest. In this condition, there was one picture for each set that elicited the highest reward across all trials and P(*Rew* I *a_optimal_*) = 70%. The original study was interested in the influence of autistic traits for reinforcement learning. For comparison with other datasets, we excluded 4 subjects who scored at a (sub-)clinical level on the autism scales used in the original study as well as 6 subjects who scored below chance level. As a result, all six models were fitted on *N* = 38 subjects for all trials (*Trials* = 90).

#### Online (Reversal 100% reward)

We collected an additional dataset to obtain the same range for P(*Rew* I a*_optimal_*) across studies in the Reversal group as in the Stable group. Jspsych-6 (de Leeuw, 2015) was used to program the task and prolific (www.prolific.co) was used to recruit 50 subjects. One subject was excluded due to chance level performance (*N* = 49). Since this dataset was not published before, we provide a more elaborate description.

As in previous work (Davidow et al., 2016; Xia et al., 2021), subjects had to direct a butterfly to its preferred flower. Each trial started with the presentation of a fixation cross for one second. Then, one (out of two) butterflies was presented together with two flowers. Butterflies differed in color. These stimuli were presented for 5 seconds or until response. On each trial, the same two flowers were presented. The locations of the flowers (left or right) were randomized over trials. Subjects had to select the left or right flower by respectively pressing the f- (left) or j- (right) key on their keyboard. Following response, the subjects received feedback for 1 second in the form of +10 or +0 points.

Since this dataset belongs to the Reversal environment, the preference of the butterflies would change during the experiment. Specifically, over the time course of 220 trials, there were 10 reversals. Rule reversals were drawn from the uniform distribution with limits 15 and 25. Notice that here P(*Rew* I *a_optimal_*) = 100%. Thus, subjects always received reliable feedback and could hence very quickly detect reversals.

Subjects were informed that each butterfly preferred one flower but that this preference could change during the experiment. After a practice phase of 40 trials with a red and green butterfly and one rule reversal, subjects were presented a two-question survey to check whether they performed and understood the task correctly. Specifically, subjects were asked (1) which butterfly preferred the white flower at the end (green or red) and they were asked (2) how many times the preference of the green butterfly changed (0, 1 or 5 times). If both questions were answered correctly, subjects could continue to the experimental phase of 220 trials with 10 rule reversals. Here, a yellow and blue butterfly (but the same flowers) were used to eliminate transfer of learning from the practice phase. The mean completion time of the study was 17 minutes for which subjects received £4 and an additional reward based on performance ranging from £1.5 to £2.5.

#### Goris et al., 2021 (Reversal 90% reward)

As mentioned before, the Goris et al. (2021) dataset contained three different conditions. One of these conditions was the volatile condition. Here, the same two sets of pictures were used but the optimal picture reversed every 18 trials (on average). Additionally, P(*Rew* I *a_optimal_*) was increased to 90%. Subjects received small monetary reward based on performance. Again, we only analyzed data of subjects who started with our condition of interest. Additionally, we excluded 5 subjects who scored (sub-)clinical on the autism scales used in the original study and 7 subjects that scored below chance level. All six models were fitted on *N* = 49 subjects for *Trials* = 90.

#### Liu et al., 2022 (Reversal 85% reward)

In this dataset (Liu et al., 2022), subjects were presented on each trial with a picture of either an animal or a piece of clothing. After this picture, two gratings were presented, one with a horizontal orientation and one with vertical orientation. The locations (left or right) of the gratings were randomized across trials. Subjects had to indicate with a left or right button press, which grating matched the initially presented picture. The grating that matched each picture was reversed after either 80 or 20 trials. During the short stable periods of 20 trials, P(*Rew* I *a_optimal_*) = 90% while during the longer stable periods of 80 trials, P(*Rew* I *a_optimal_*) = 80%. Although P(*Rew* I *a_optimal_*) was not stable during the whole experiment, we believed changes were too small to categorize this dataset in the Stepwise group. Subjects did not receive monetary reward based on performance. Again, all six models were fitted for all *N* = 23 subjects on all trials (*Trials* = 320).

#### Verbeke et al., 2021 (Reversal 80% reward)

In this dataset (Verbeke, Ergo, De Loof, & Verguts, 2021), Subjects were presented either a horizontal or vertically oriented grating. Subjects had to learn which response, right (j-key) or left (f-key) matched with each grating. There were 15 reversals and the response that matched each grating was reversed every 15-45 trials. Here, P(*Rew* I *a_optimal_*) = 80% across the entire task. Subjects received a small monetary reward based on performance. All six models were fitted for all *N* = 27 subjects on all trials (*Trials* = 480).

#### Mukherjee et al., 2020 (Reversal 70% reward)

In this dataset (Mukherjee et al., 2020), subjects were presented with two fractals, one on each side of the screen. The locations of the fractals were randomized across trials. Subjects chose one of the fractals by pressing a keyboard button. After the response, feedback was presented in the form of a coin (reward) or a red dot (absence of reward). Here, P(*Rew* I *aoptimal*) = 70% across the entire task. Subjects received a small monetary reward based on performance. In the original study, there was a reward and a punishment condition. Additionally, two groups of subjects were used. One group with a major depressive disorder and a control group. For comparison with the other datasets, we only used data from the reward condition for the control group. Additionally, 8 subjects were excluded due to below chance level performance. As a result, all six models were fitted on the data of *N* = 56 subjects for all trials (*Trials* = 90).

#### Cohen et al., 2007 (Stepwise)

In this dataset, the subject was presented with two grey squares, one on each side of the screen. Subjects simply had to press either a right or left button on their keyboard. After the response, feedback was presented in the form of +10 or -10 cents. Every 80-150 trials, the probability of reward given a particular button press changed (but, unlike the reversal environment, did not typically revert to its previous level; see Figure 1B for illustration). Here, P(*Rew* I *a_optimal_*) could be either 75% or 50%. Subjects received small monetary reward based on performance. Two subjects were removed from the analyses due to chance level accuracy. All six models were fitted for the remaining *N* = 15 subjects on all trials (*Trials* = 1200).

#### Hein et al., 2021 (Stepwise)

In this dataset, subjects were presented with two fractals, one on each side of the screen. The locations of the fractals were randomized across trials. By making a button press on the keyboard, subjects needed to select one of the fractals. After the response, subjects received feedback (win 5 pence or lose 5 pence). Every 26-38 trials the probability of reward given a particular fractal changed. Here, P(*Rew* I *a_optimal_*) could be 90%, 70% or 50%. Subjects received small monetary reward based on performance. In the original study, subjects were split into two groups based on their score for an anxiety trait questionnaire. For comparison with the other datasets, we only used data from the control (not anxious) group. As a result, all six models were fitted on the data of *N* = 20 subjects for *Trials* = 400.

### Analyses

#### Model simulations and fitting

Parameter recovery and model recovery simulations were initially carried out for each model. As we argued in previous work (Beeckmans et al., 2023), using plausible value distributions for parameter recovery is preferable to using the whole range of possible values. Therefore, Recovery simulations were informed by our previous work (Verbeke, Ergo, De Loof, & Verguts, 2021) . Here, a subset (3/6) of models (see Fig. 1A) were fitted on a reversal learning task (480 trials with 15 reversals). Based on this previous study (Verbeke, Ergo, De Loof, & Verguts, 2021), we determined plausible value distributions for each parameter in each model. From these distributions, 540 values for each parameter were drawn and used to simulate 540 datasets on the design of the reversal learning task. This resulted in 3240 (540 x 6 models) simulated datasets on the reversal learning task. We compared the true distributions of values for each parameter to the estimated distributions and computed the Pearson correlation coefficient between the true and estimated parameters.

Importantly, since the main goal of the current study is to identify which computational strategy is most likely to underlie a specific behavioral dataset, we also performed model recovery analyses (Wilson & Collins, 2019). Here, we used weighted Akaike information criterion values (see below) to evaluate which model was most likely to have generated a particular dataset. Ideally, one would want the Akaike weights to be high only for the model that generated that dataset.

We next aimed to investigate which model (see Fig. 1A) was able to perform with the highest accuracy in each environment. For this purpose, we extracted for each subject the experimental design (i.e., stimulus sequence, rewarded response, …). We used these designs to estimate the parameters that optimized total reward for each model. For optimization, the differential evolution method was used as implemented by the SciPy (version 1.4.1) package in Python (version 3.7.6). The optimized parameters were used to simulate each model on each dataset. Here, accuracy was computed as the proportion with which the optimal action was selected (P(a_optimal_)). For easy comparison, across models and datasets, we constructed a novel weighted accuracy measure (wACC). Here, accuracy was averaged over simulations and transformed to a weighted measure of accuracy. Specifically,

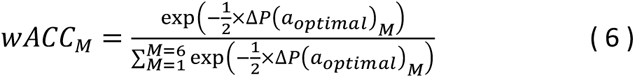

in which

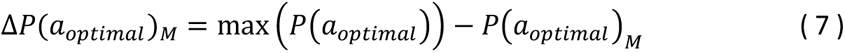

Hence, the wACC represents how strongly the accuracy of model *M* deviated from the most optimal (max) model. Here, better performing models have a higher wACC and the wACC across all models always sums to 1.

The wACC was computed twice. First, wACC was computed for each cognitive model in comparison to the other 5 cognitive models. Second, wACC was computed for the six cognitive models but also including two benchmarks. As mentioned before, one benchmark was the GRU network. Another benchmark was the performance of the subjects in each dataset.

As for the cognitive models, the GRU network was trained to optimize total reward. For this purpose, it received the rewarding response at each trial as target. The network was trained until the cross-entropy loss remained stable (i.e., decreased less than .001) for 10 consecutive episodes. Here, an episode consisted of all trials in an experimental design and weights were updated every 10 trials. The total number of episodes never exceeded 70. After training, weights were fixed, and the model was tested again on the same experimental design. Again, accuracy was computed as P(*a_optimal_*).

After obtaining the best performing model, we next aimed to investigate which model fitted best with human behavioral data. For this purpose, parameter estimation was performed for each model of Fig. 1A on each dataset in Table 1. Parameter estimation was implemented by minimizing the negative loglikelihood (LL) via the differential evolution method using the SciPy (version 1.4.1) package in Python (version 3.7.6).

Again, we used weighted measures for easier comparison across models (Wagenmakers & Farrell, 2004).

Here,

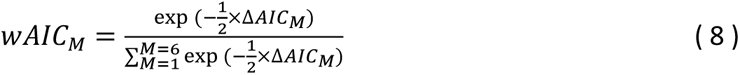

for which

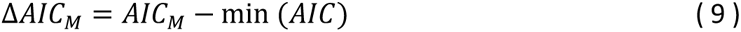

in which AIC is the Akaike information criterion (AIC). This AIC measure adds a small penalty (2 x *k*) for the number of parameters (*k*) to the negative loglikelihood. Again, the weighted measure represents how strongly the fit of model *M* deviated from the most optimal (min) model. Here, models that fit better have a higher wAIC and the wAIC across all models always sums to 1. To explore how the penalty for number of parameters influences our results, we also computed weighted measures of fit based on the loglikelihood (no penalty) and on the Bayesian Information criterion (BIC; penalty of ln(*Trials*) x *k*).

Each model was simulated and fitted for each individual subject. Based on the simulations, we extracted the wACC for each model in each environment. Based on results of the fitting procedure, wAIC (but also wLL and wBIC) values were computed for each subject in each dataset.

Since the wAIC only informs about how well a model fits compared to the other models in the comparison, we also provide a measure of normalized fit which allows to determine the goodness of fit for each model in an absolute manner. Specifically, this measure provides a percentage between 0 and 100% of how well the model fits the data. This measure is formalized as

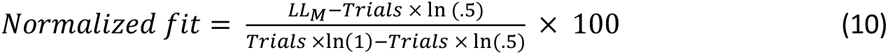

Here, *LL_M_* represents the optimized loglikelihood of the model (M) on the behavioral data. This is evaluated compared to a lower bound of Trials x ln(.5), reflecting a situation where the model provides a .5 likelihood to each response on each trial, and an upper bound of Trials x ln(1) which always equals 0 and reflects a situation where the model was able to predict every response with a likelihood of 1. Hence, the normalized fit represents how well the model fits the data in between this lower and upper bound.

#### Linear mixed models

To provide some general insight in the influence of the environment and P(*Rew*|*a_optimal_*) on behavioral performance, a linear mixed model regression was performed with averaged accuracy for each human subject in each dataset as dependent variable. Environment (stable, reversal or stepwise) and P(*Rew*|*a_optimal_*) were used as fixed independent variables. Dataset was included as random intercept.

Additionally, to further evaluate the added value of each separate hierarchical model extension, we performed two linear mixed model regressions with either wACC or wAIC as dependent variable and environment (Stable, Reversal or Stepwise), adaptive learning rate (yes or no), multiple rule sets (yes or no) and hierarchical learning (yes or no) as fixed independent variables. Dataset was again included in the analyses as a random intercept.

## Results

### Descriptive statistics of the datasets

As described in the Methods, we collected ten empirical datasets across three types of environments (see Figure 1B). Specific features of each dataset are summarized in Table 1 (see Methods). Fig. 2A provides some descriptive analyses for each dataset. Accuracy is defined as the proportion of optimal actions (i.e., actions that would elicit reward if feedback was deterministic; P(*a_optimal_*)). The effect of environment on overall accuracy was not statistically significant (χ^2^(2, *N* = 407) = 4.572, *p* = .102) but accuracy did decrease significantly (χ^2^(2, *N* = 407) = 13.592, *p* = .0002) for lower values of P(*Rew*|*a_optimal_*). This is observed as a decreasing accuracy trend from left to right in each grey box of Fig. 2A. Notably, the Goris/Volatile dataset, seems to be a strong outlier in this trend. While P(*Rew*|*a_optimal_*) = 90% in this dataset, accuracy is lower than for Reversal datasets where P(*Rew*|*a_optimal_*) is 85 or 80%. This difference in comparison to other Reversal datasets might be due to the fact that rule reversals occurred faster on average than in the other datasets (see Table 1).

**Figure 2.**
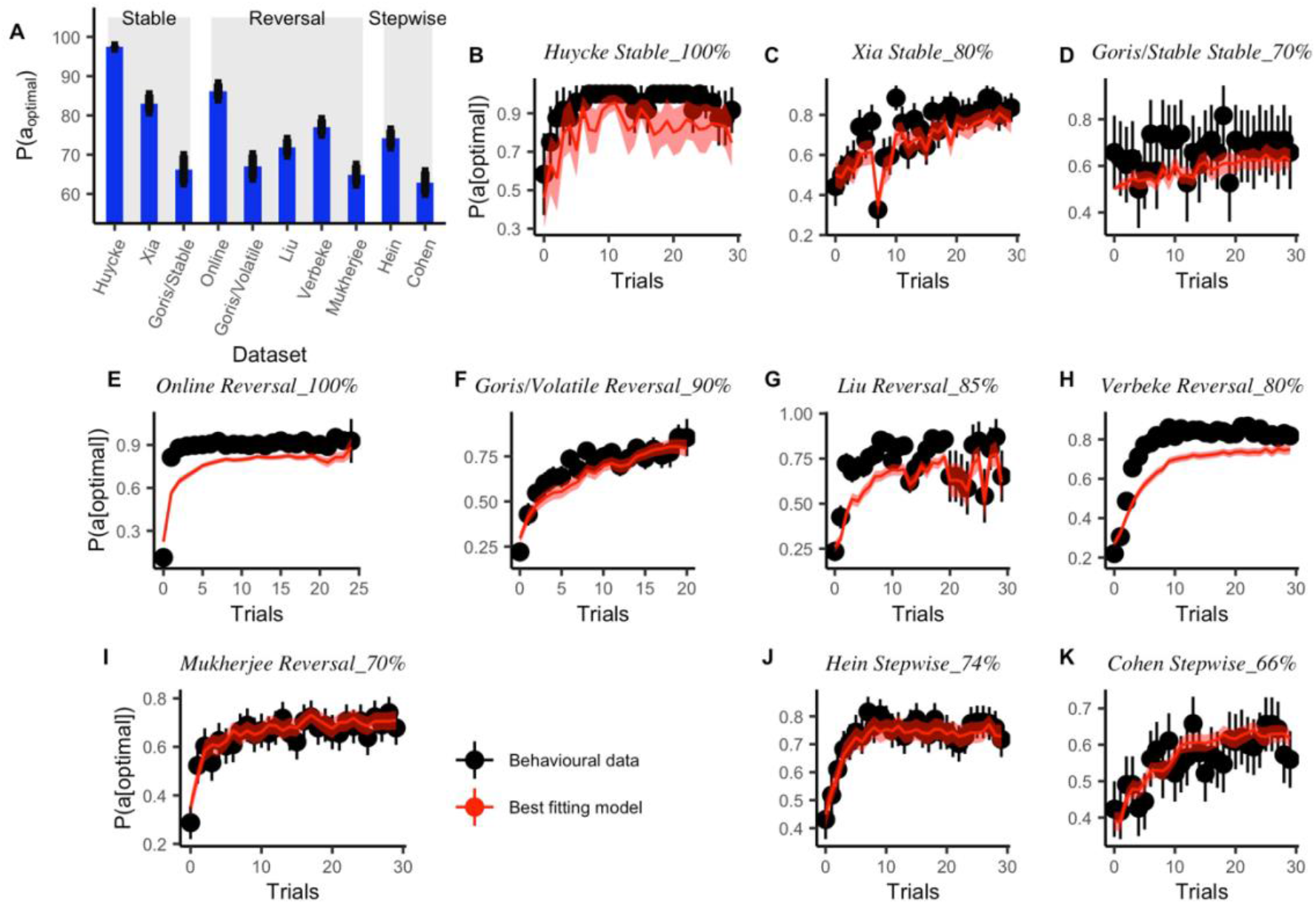
Datasets. *A:* Overall task accuracy in each dataset. Blue bars indicate the average accuracy (defined as proportion of optimal actions; P(*a_optimal_*)) across all subjects for each dataset. Black lines represent the 95% confidence interval. Datasets are grouped for different environments (grey boxes) and sorted for P(*Rew*|*a_optimal_*) in descending order. *B-K:* Learning curves for each of the 10 datasets. Here, trial 0 is the moment where reward contingencies changed, or in the case of Stable environments, the first trial of the experiment. Black dots and bars indicate behavioral accuracy of the human subjects and the 95% confidence interval. The red line represents the fitted learning curve for the model with overall best fit (see also Fig. 7B).

Fig. 2B-K also illustrate the behavioral learning curves for each dataset respectively. Here, the change in accuracy is shown for trials since a rule switch (for the Reversal and Stepwise environments) or since the start of the tasks (for the Stable environment). The performance of the human subjects is shown by the black dots. The red lines illustrate the fitted learning curves for the best fitting model (in terms of wAIC; see also Fig. 7B).

### Model recovery

To check the validity of the models, we performed simulations to determine the parameter and model recovery. For these simulations, we used parameter distributions from previous work (Verbeke, Ergo, De Loof, & Verguts, 2021). Fig. 3 shows the true parameter distributions that were used for simulations and the parameter distributions that were estimated based on this simulated data. As can be observed, these distributions are largely similar for most of the parameter values across all models. Notably, there is a discrepancy between the true and estimated distributions for the learning rate in models with an adaptive learning rate (ALR, Sets_ALR and Full model). Indeed, this value is much harder to estimate in these models than in the other models, because the learning rate in models with an adaptive learning rate only represents the starting value of the learning rate on the very first trial. After this first trial, the learning rate is changing with a rate determined by the Hybrid parameter. The rate at which the learning rate is changing (i.e., the Hybrid parameter) is stable over trials and is thus easier to estimate. Importantly, this also means that the (initial) learning rate in these models is of much less importance for model behavior than learning rate is in the other models.

**Figure 3.**
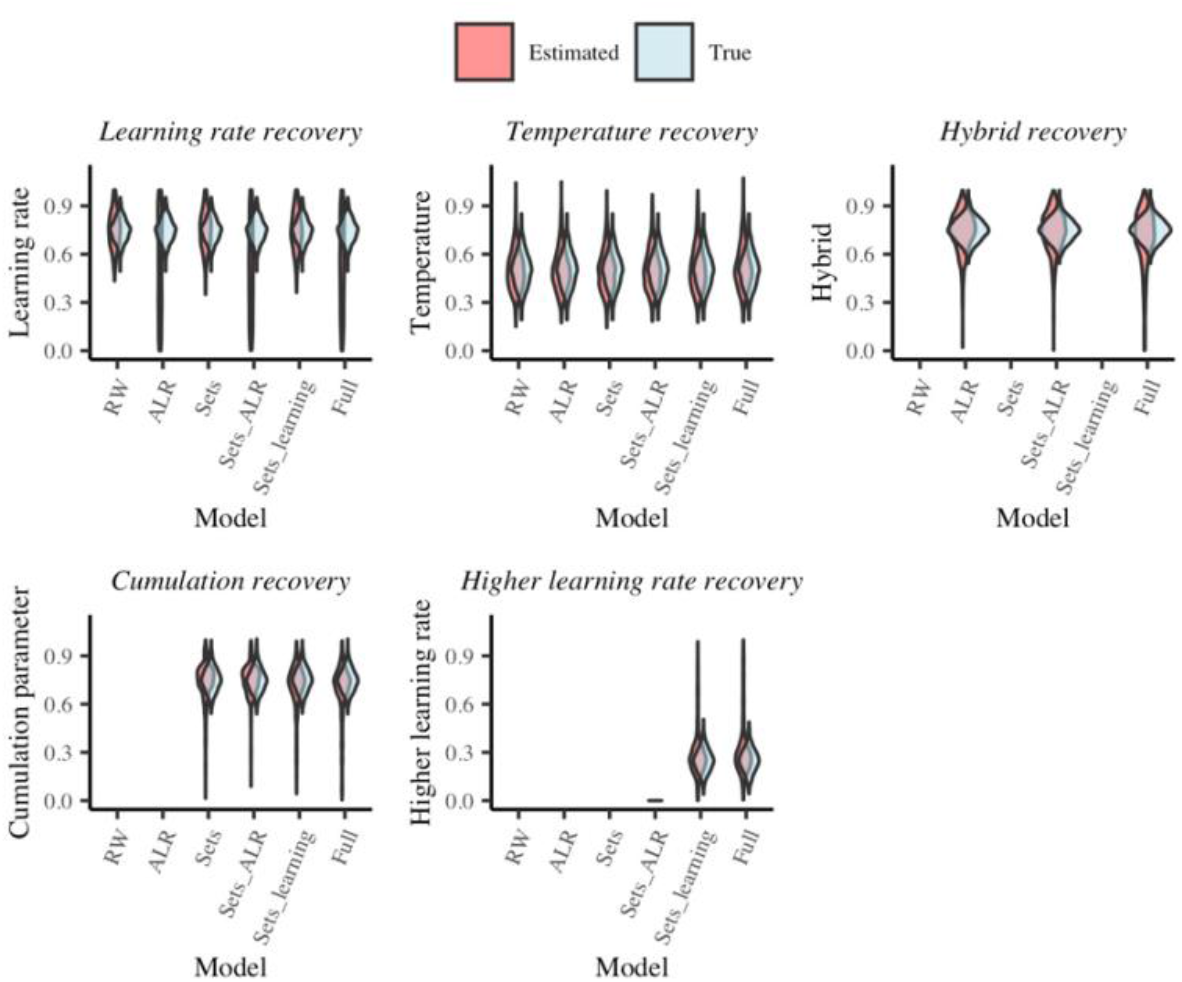
True and estimated distributions for parameter values in parameter recovery analyses. If there is no distribution in the panel it is because this parameter is absent in that particular model.

The Pearson correlation coefficient between the true and estimated parameters is reported in Fig. 4A. Here, white cells represent parameters that are not present in that model. As expected from Fig. 3, the correlation for the (initial) learning rate is indeed very low for models with an adaptive learning rate. For other parameters, the correlations are reasonably high. Apart from the learning rate in models with an adaptive learning rate, the lowest observed correlation value is .293 which is still highly significant (p < .0001).

**Figure 4.**
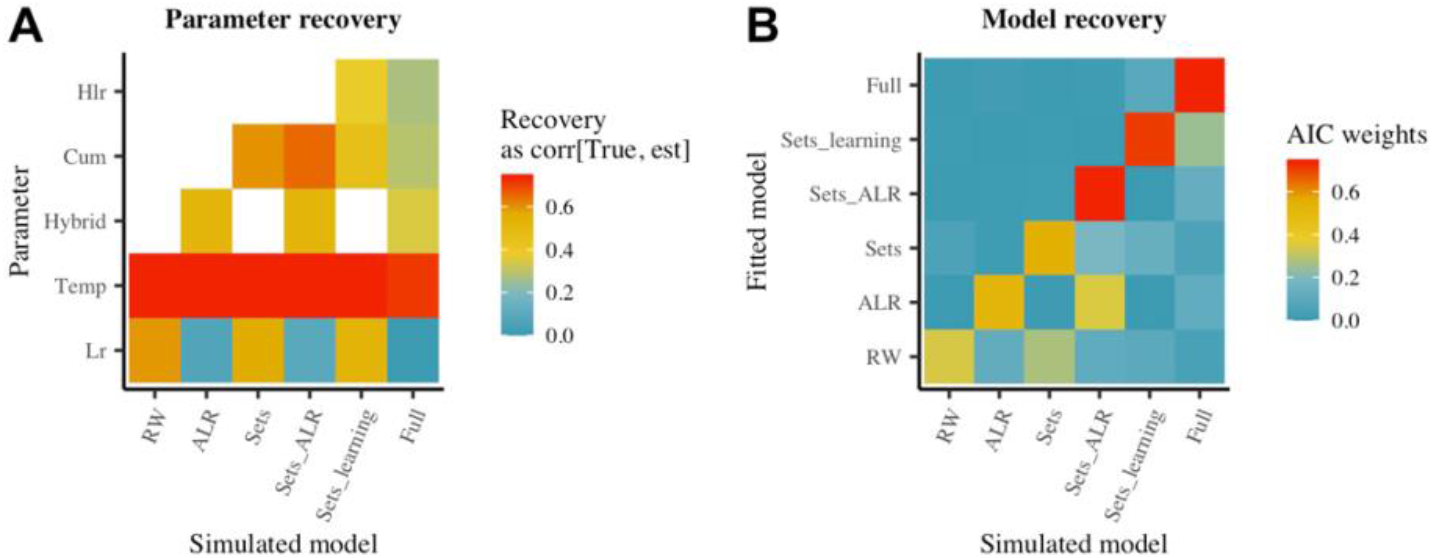
Recovery results. *A: Parameter recovery*. Shown as the Pearson correlation between the true and estimated parameters. Here, white cells indicate parameters that are absent for that particular model. *B: Model recovery results*. Here, the color represents the weighted (AIC) evidence for a particular model.

Nevertheless, the main goal of the current study is to identify what computational strategy is most likely to be underlying a specific behavioral dataset. As shown in Fig. 4B, we can indeed clearly distinguish which model was responsible for a particular dataset. The more parameters in the model, the more distinct behavioral dataset it produces and hence the stronger the evidence for that particular model.

### Best performing model depends on the environment

First, model performance was evaluated in each environment. For comparison, we use a weighted accuracy measure (wACC; see Methods for details). Hence, each cell in Fig. 5 represents the relative accuracy for a specific model (columns) in a specific environment (rows) compared to the accuracy of other models in that environment (sum of each row is 1). Fig. 5A shows results if one compares only the cognitive models while Fig. 5B adds two benchmarks in this comparison. One benchmark is the GRU network and the other benchmark is the accuracy of the human subjects (see also Fig. 2A).

**Figure 5.**
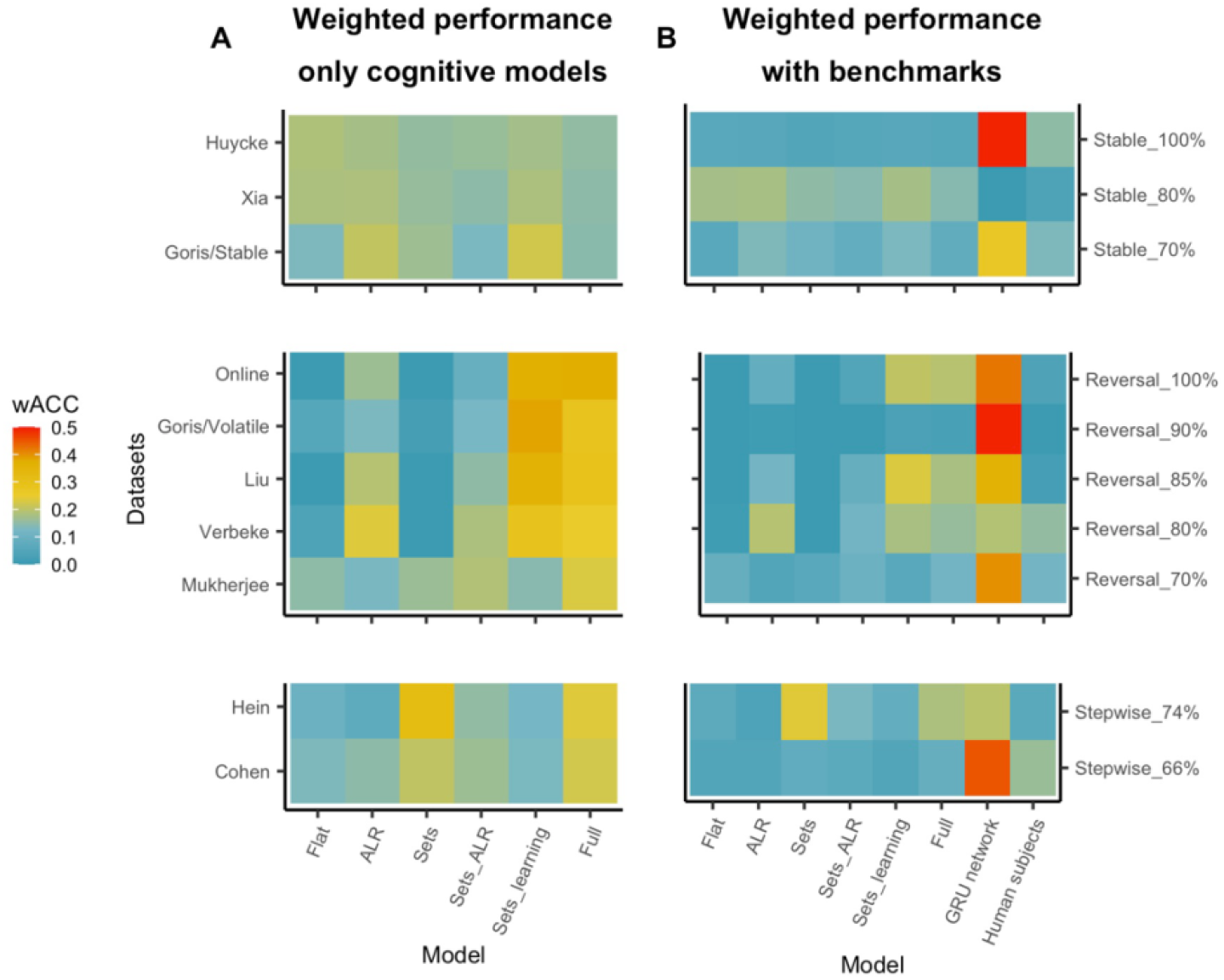
Model performance. A weighted measure of model performance (wACC; see Methods) is shown for each model and each dataset. Model performance is weighted to allow comparison across models, each row sums to 1. Datasets are again grouped per environment and sorted for P(*Rew*|*a_optima_*_l_). *Row A* demonstrates a comparison of only the 6 cognitive models and *Row B* includes two benchmarks in the comparison: a GRU network and the performance of the human subjects.

A direct comparison of the cognitive models (Fig. 5A) demonstrates that for the Stable environment, there is no strong difference in performance across models. Hence, in this environment there is no benefit in adding parameters to the Flat model. In the Reversal environments, there is a clear benefit in performance for additional model parameters. Here, best performance is observed for the Sets_learning and Full models. Both these models implement multiple rule sets and hierarchical learning. The Full model also adds an adaptive learning rate. Also in the Stepwise environment a benefit in performance is observed for hierarchical model extensions. Here, the Sets and Full models outperform the other cognitive models. Both these models add multiple rule sets to the Flat model.

When adding the two benchmarks to the comparison (Fig. 5B), it is clear that the GRU network outperforms the cognitive models. In this sense, the cognitive models cannot reach optimal performance. Nevertheless, in most datasets, the cognitive models perform within reasonable range from the GRU network. The mean difference in accuracy between the best cognitive model and the GRU network was 8.078 % (SD = 4.22across all datasets. Moreover, in almost all datasets the human subjects performed worse than the cognitive models. Notably, this difference in performance between humans and the best cognitive model (Mean = 8.590%; SD = 3.835). was smaller than between the human subjects and the GRU network (Mean = 9.366%, SD = 5.330). In conclusion, the cognitive models are not optimal but they do a good job (better than the GRU model) of mimicking human performance in this task.

### GRU network switches between modules

Since the GRU network outperforms the cognitive models, we aimed to investigate how the neural network deviates from these cognitive models. Specifically, we investigated network activation for our simulations on the Mukherjee dataset (Reversal environment). This dataset consisted of 90 trials divided in three blocks of 30 trials following an ABA pattern. Hence, subjects had to learn rule A, then after 30 trials switch to rule B and switch back to rule A for the last 30 trials. Figure 6A shows distributions over 23 simulations (one for each participant design) for the representational distance (Euclidian distance of activation patterns) between the first and second occurrence of task rule A (in blue) and between the first occurrence of task rule A and the first occurrence of task rule B (in red). This demonstrates that the network activation is very similar during the first and second occurrence of the same task rule but very different for the first occurrence of different task rules. This is not trivial since a recurrent network integrates activity over time and hence network activation at one point in time is typically very similar to activation at the surrounding time points. In contrast, the first and second occurrence of task rule A are much more separated in time than the first occurrence of task rule A and the first occurrence of task rule B. This suggests indeed that, as implemented in the cognitive models with multiple rule sets, the network might construct two clearly separate representations and perform an abrupt switch once the rule has changed. The better performance might in that case be caused by a better switching strategy (better hierarchical learning).

**Figure 6.**
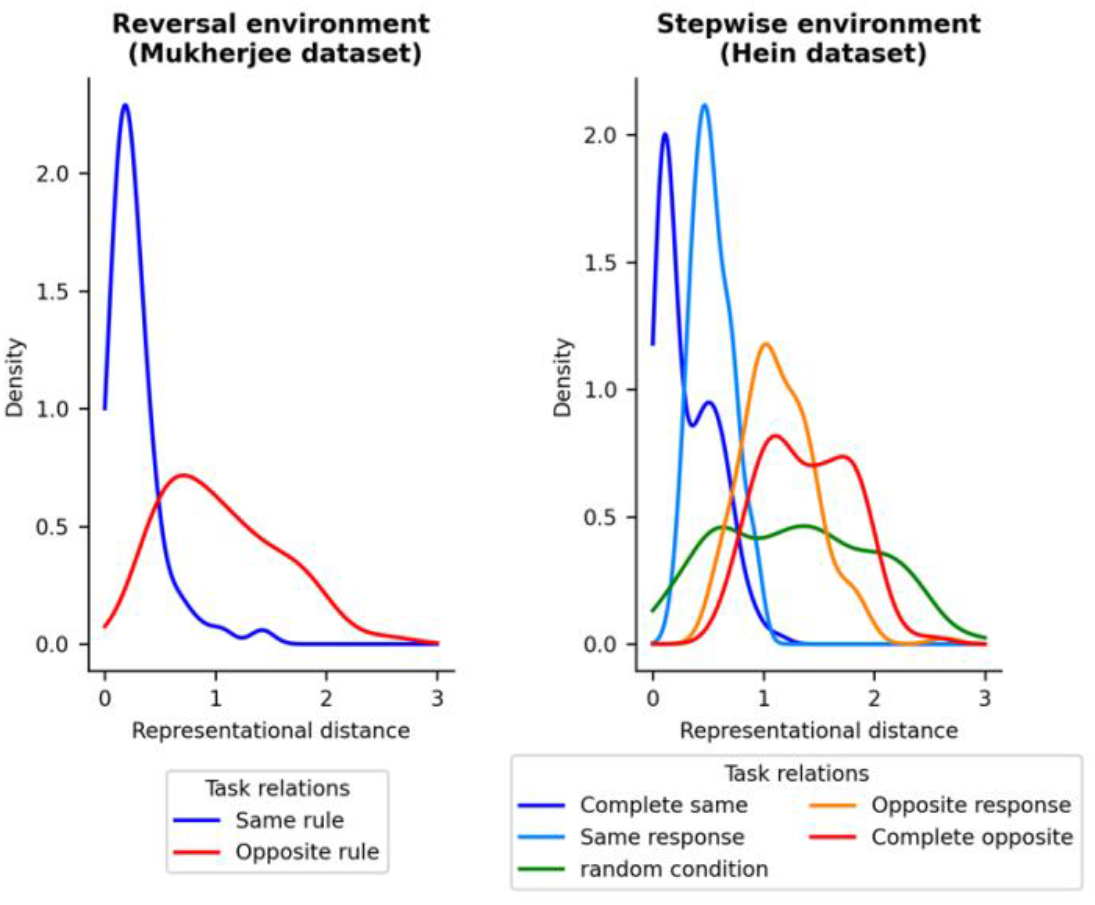
Representations in the GRU network. For each dataset, one simulation of the GRU network is performed for each participant in that dataset. Distributions depict the Euclidean distance in activation patterns at the hidden layer between two types of task blocks (for multiple simulations). Here, task blocks could be completely the same, meaning that they have the same reward probability and the same optimal action (*Same rule* or *Complete same* in legend), or they could be completely opposite in the sense that the reward probability for both actions has been inverted and hence the identity of the optimal action is different, but the same reward probability applies for that other action (*Opposite rule* or *Complete opposite* in legend). In the Stepwise environment, it is also possible that for two task blocks the identity of the optimal action remains the same, but the reward probability is different (*Same response* in legend). For the *Opposite response* task rules, both the reward probability and the optimal action are different. Because the 50% reward probability is a random condition and there is no optimal action, a different distribution is made that compares this task rule to all other task rules. *A:* shows representational distance for the Mukherjee dataset in the Reversal environment. *B:* shows representational distance for the Hein dataset in the Stepwise environment.

**Figure 7.**
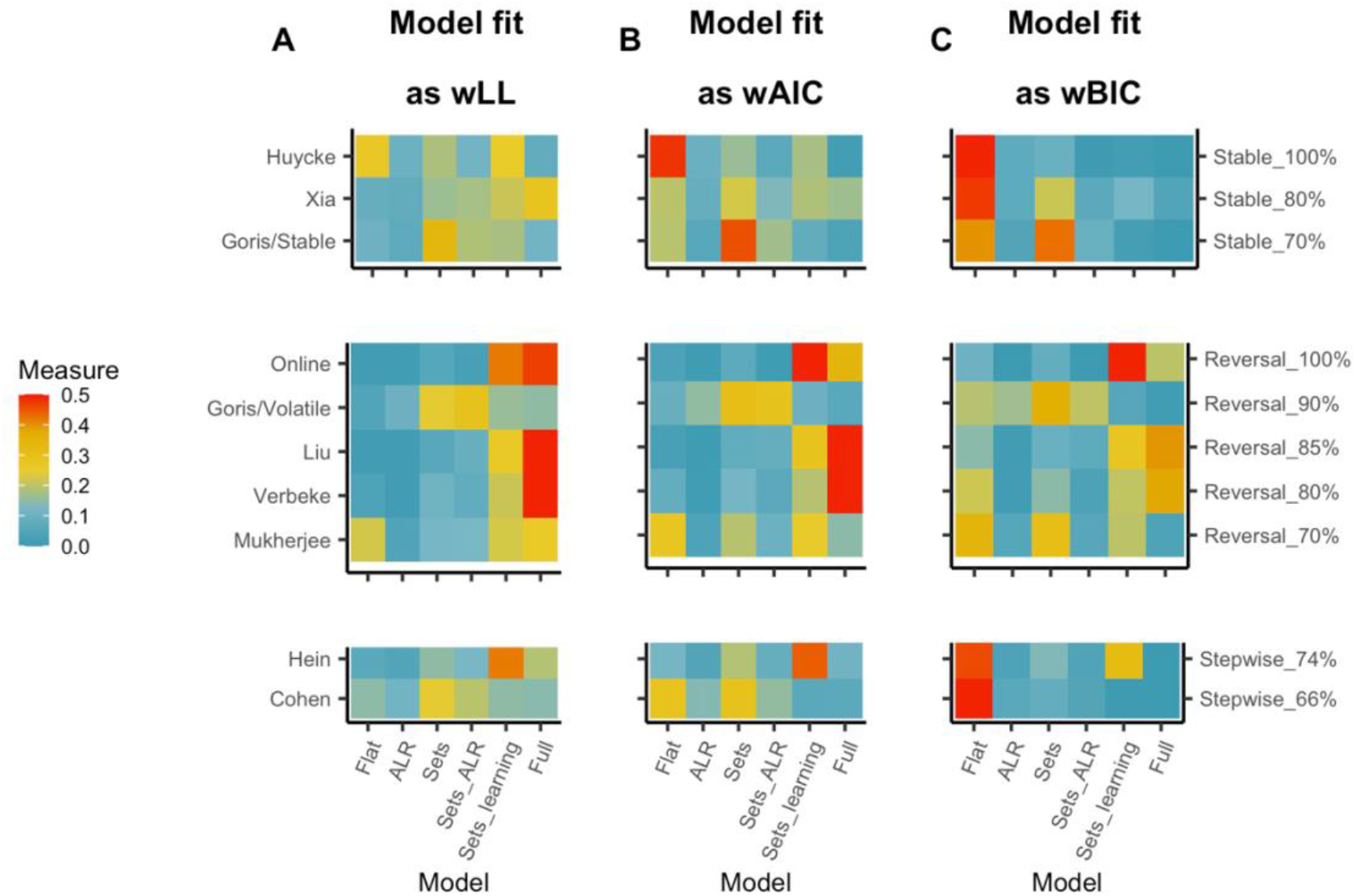
Model fit. Weighted measures of fit are shown for each model on each behavioral dataset. Model evidence is weighted to allow comparison across models, each row sums to 1. Datasets are again grouped per environment and sorted for P(*Rew*|*a_optima_*_l_). *Column A* demonstrates a comparison of loglikelihood which does not include a penalty for the number of parameters. *Column B* demonstrates a comparison of AIC which includes a small penalty for additional parameters. *Column C* demonstrates a comparison of BIC which severely penalizes additional parameters.

Similar analyses were performed for the Stepwise environment. Figure 6B shows results for the Hein dataset. In this task, there were 5 levels of P(*Rew*|*a* = 1). Here, P(*Rew*|*a* = 1) could be 90, 70, 50, 30 or 10%. In the trials where P(*Rew*|*a* = 1) < 50%, this of course means that the identity of the optimal action has changed and hence P(*Rew*|*a* = 2) > 50%. In the P(*Rew* | *a* = 1) = 50 %, also P(*Rew* | *a* = 2) = 50 %. Hence, in this condition it is perfectly optimal to respond at random. In this environment, the cognitive models with multiple rule sets would ignore the difference between the P(*Rew*|*a_optimal_*) = 90 and 70% condition and just construct two rule sets: One for when P(*Rew*|*a* = 1) > 50%, and one for when P(*Rew*|*a* = 2) > 50%. In the P(*Rew* | *a* = 1) = 50 % the model could then select one of these rule sets at random. However, Figure 6B demonstrates that the GRU network indeed constructs 5 rule sets and hence demonstrates different activation patterns when the optimal response is still the same, but the reward probability is different (light blue distribution). This suggests an alternative approach in which both the identity of the optimal action and the reward probability are taken into account.

### More complex models fit human behavior better in more complex environments

Next, model fit to empirical data was evaluated. As illustrated in Fig. 2B-K, models were able to adequately describe averaged learning curves of the human subjects. Furthermore, Table 2 reports Normalized fit (see Methods) for all models on each dataset. A more detailed insight in model dynamics is given in the Supplementary materials (trial-by-trial comparisons between responses and model predictions; Figures S2 and S3).

**Table 2.** Normalized fit.

|  | *Flat* | *ALR* | *Sets* | *Sets_ALR* | *Sets_Learning* | *Full* |
| --- | --- | --- | --- | --- | --- | --- |
| *Goris/Stable* | 20.98 | 17.27 | 25.73 | 19.10 | 22.38 | 18.71 |
| *Xia* | 56.92 | 56.67 | 58.95 | 59.17 | 59.77 | 60.3 |
| *Huycke* | 90.00 | 87.96 | 89.62 | 87.75 | 89.77 | 87.31 |
| *Mukherjee* | 26.18 | 35.08 | 38.60 | 38.35 | 40.93 | 40.76 |
| *Verbeke* | 23.36 | 22.48 | 26.91 | 26.12 | 27.82 | 29.13 |
| *Liu* | 30.75 | 30.42 | 32.45 | 32.36 | 35.81 | 37.81 |
| *Goris/Volatile* | 38.45 | 41.39 | 48.45 | 44.29 | 43.10 | 42.50 |
| *Online* | 29.27 | 26.59 | 35.82 | 34.28 | 49.97 | 49.87 |
| *Hein* | 45.24 | 44.79 | 46.03 | 45.48 | 47.14 | 46.18 |
| *Cohen* | 13.66 | 13.35 | 13.77 | 13.63 | 13.68 | 13.17 |

Notes. Each cell provides the normalized fit for a particular model on a particular dataset (see Methods on how normalized fit is computed).

To efficiently compare models, Figure 7 shows three weighted measures of fit: weighted log-likelihood (Fig. 7A), weighted AIC (Fig. 7B) and weighted BIC (Fig. 7C). Hence, empirical support for each model is evaluated by using different penalties for the number of parameters in the model. While the log-likelihood uses no penalty for adding parameters, the AIC implements a mild penalty, and the BIC measure implements a stronger penalty (see also Methods for details). Hence, each cell in Fig. 7 represents the relative goodness of fit for a specific model (columns) in a specific dataset (rows) for a specific measure (panels A, B or C).

Implementing stronger penalties for the number of parameters naturally shifts the weighted measures of fit to the simpler models. However, across all three measures of fit, it is clear that more hierarchically complex models fit better in the Reversal environment than in the Stable environments.

Similar to the results of model performance (Fig. 5A), weighted loglikelihood evidence in the Stable environment is similar across all models. As a result, the simplest Flat model receives the highest weight when a penalty for the number of parameters is applied. Note that this is less clear in the Goris/Stable dataset. Unless one implements a severe parameter penalty, such as in BIC (Fig. 7C), a better fit is observed for the Sets model in this dataset. This suggests a strategy where subjects switch their behavior when they did not get a reward for a certain number of consecutive trials. This number of trials before a behavioral switch occurs, is determined by the Cumulation parameter (See Methods and Fig. 1A). Note that the Goris/Stable dataset also has the lowest P(Rew|*a_optimal_*) = 70%. Hence, it is possible that, although *a_optimal_* is stable, subjects never fully converged to that action. Instead, they constantly test and alternate between the two alternative hypotheses on *a_optimal_*.

In the Reversal environment the results are consistent irrespective of the penalty for the number of parameters that is applied. Also here, the pattern is strikingly similar to the results of model performance (Fig. 5A). The fit to behavioral data is best for the Sets_learning and Full model. Both these models implement multiple rule sets and hierarchical learning, but the Full model also adds an adaptive learning rate. Notably, the Mukherjee and Goris/Volatile dataset also demonstrate considerable evidence for the Sets model. According to the model performance results (Fig. 5A) this is not the best strategy to use but accuracy is also lowest for these datasets (see Fig. 2A). In the Mukherjee dataset this lower accuracy is probably due to the higher volatility with P(*Rew*|*a_optimal_*) = 70%. In the Goris/Volatile dataset this lower accuracy is probably due to the low number of trials between every reversal (18 trials on average; see Table 1). Hence, it is possible that in such high volatile environments, subjects prefer a simpler heuristic/model than in the less noisy environments.

For the Stepwise environment, evidence depends on the penalty that is applied. With a severe penalty for the number of parameters (Fig. 7C), the Flat model fits best in both datasets. However, in the other measures with a lower penalty, the Sets model fits best in the Cohen dataset, and the Sets_learning model fits best in the Hein dataset. This suggests that also in the Stepwise environment there is a potential benefit for multiple rule sets in model fit (Fig. 7) and, as noted before, also in model performance (Fig. 5A)

In sum, results suggest that overall humans adaptively select the model that results in the best performance. Which model performs or fits best differs across environments: In the Stable environment, the Flat model is most suited. When the environment becomes more complex, there is also more evidence for more complex models, especially for the models that implement multiple rule sets.

### Humans adaptively combine hierarchical extensions

A key point of the current study was to investigate the added value of each individual hierarchical extension in each of the three environments. To shed further light on this, the left column of Fig. 8 (panels A, C and E) presents the wACC of model simulations averaged for each combination of environment and hierarchical extension. We also carried out linear mixed model analysis with environment and extension as (categorical) factors and wACC as dependent variable. A summary of the statistical tests is given in Table 3, but we discuss the results in more detail below.

**Figure 8.**
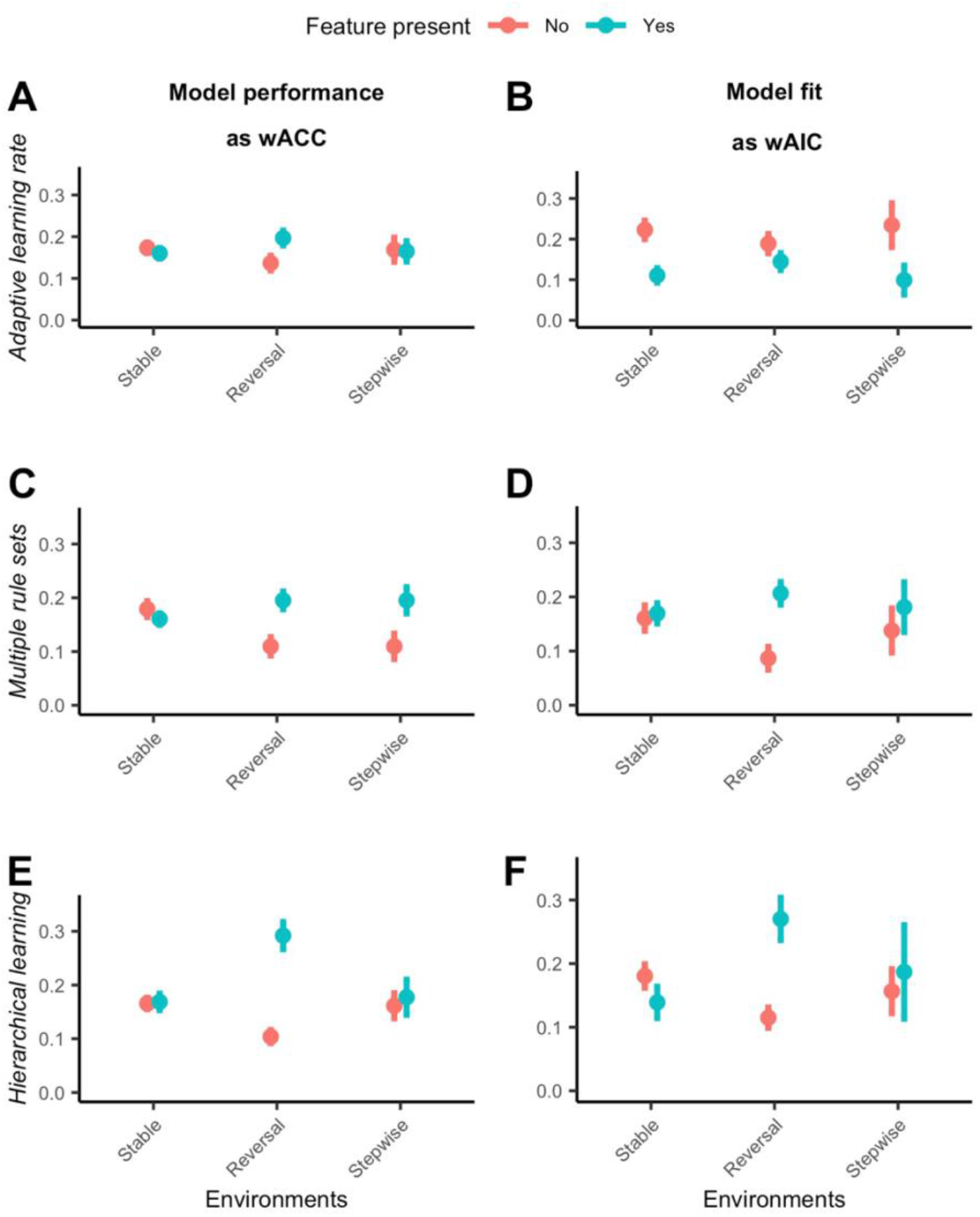
Hierarchical model extensions in each environment. Each datapoint reflects the mean (wACC or wAIC) over all models with one particular feature (e.g., the ALR, Sets_ALR and Full model for the adaptive learning feature) in a particular environment. *Column 1(A, C, E)* illustrates the influence of each extension on model performance (as wACC on y-axis) across environments. *Column 2(B, D, F)* illustrates influence of each extension on model fit (as wAIC on y-axis) for the behavioral datasets in each environment.

**Table 3.**
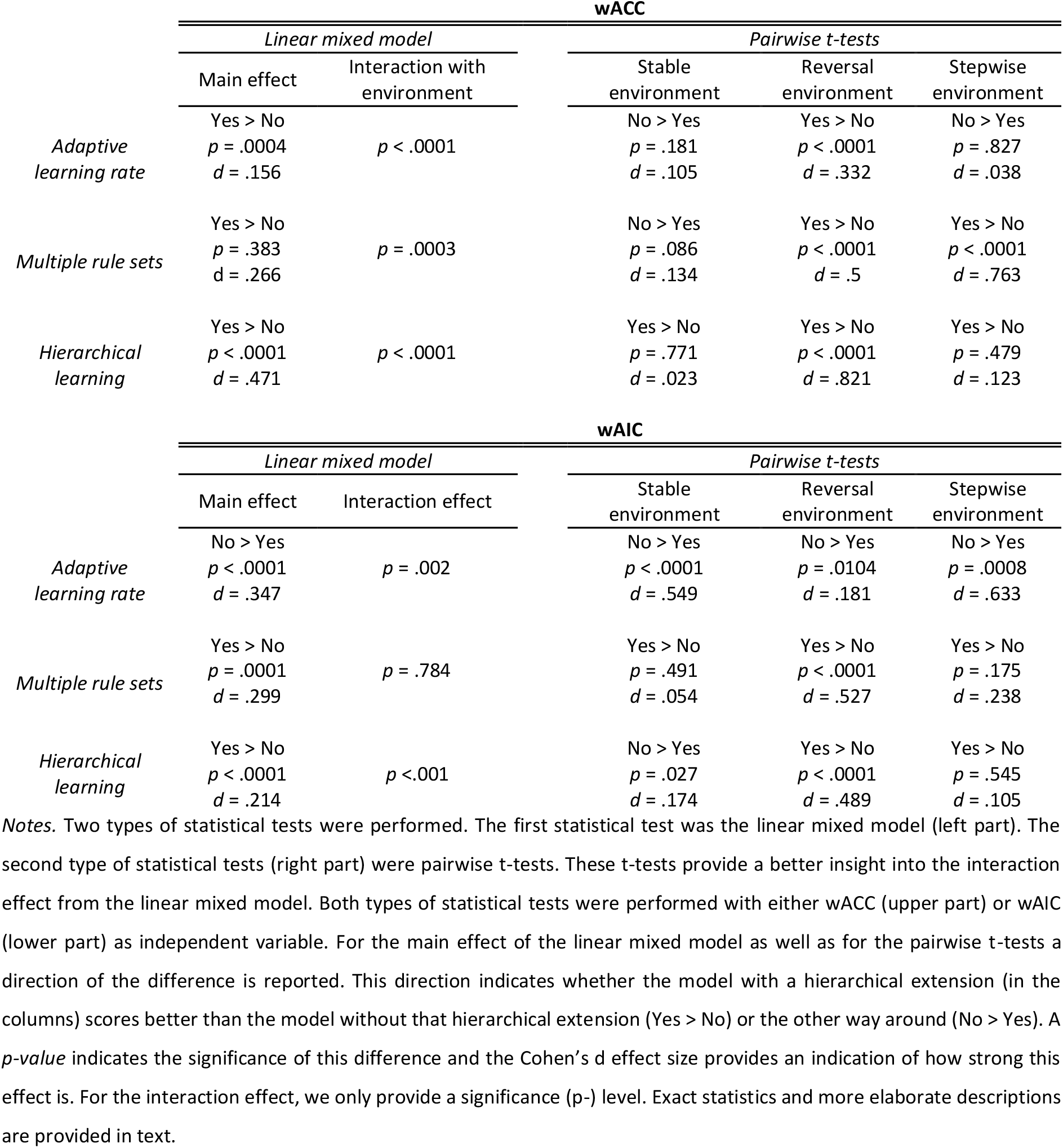
Statistics.

For the adaptive learning rate feature, a main effect was found (χ^2^(1, *N* = 407) = 12.24, *p* = .0004) showing that the models performed significantly better overall if they used an adaptive learning rate (*mean wACC* = .179) than when they did not (*mean wACC* = .154). However, as shown in Fig. 8A, this effect is mainly driven by the Reversal environment. Indeed, the linear mixed model reported a significant interaction between the adaptive learning rate and the environment (χ^2^(2, *N* = 407) = 25.606, *p* < .0001), showing better performance (*t*(205) = 4.749, *p* <.0001) with adaptive learning rate (*mean wACC* = .197) than without adaptive learning rate (*mean wACC* = .137) in the Reversal environment but no significant differences in the Stable (*t*(165) = -1.344, *p* = .181) or Stepwise (*t*(34) = -.220, *p* = .827) environments. Furthermore, there was a significant three-way interaction between environment, adaptive learning rate and hierarchical learning (χ^2^(2, *N* = 407) = 25.1620, *p* < .0001), indicating that the benefit of an adaptive learning rate in the Reversal environment is mainly driven by the implementation of the adaptive learning rate in models that also implement hierarchical learning (i.e., the Full model).

For multiple rule sets, no significant main effect was observed (χ^2^(1, *N* = 407) = .760, *p* = .383) in the mixed linear model. Nevertheless, a t-test (*t*(406) = 5.364, *p* < .0001) did indicate that models performed better overall with multiple rule sets (*mean wACC* = .181) than without (*mean wACC* = .137). This discrepancy between the linear mixed model and t-test occurs because the linear mixed model also includes the hierarchical learning feature which is dependent on the multiple rule sets and hence explains largely the same variance. Investigating the performance benefits of the multiple rule sets feature in more detail, a significant interaction with environment is observed (χ^2^(2, *N* = 407) = 16.134, *p* = .0003). More specifically, as illustrated in Fig. 8C, there is no significant benefit in the Stable environment (t(165) = -1.725, p = .086) but there is a strong benefit in the Reversal environment (*t*(205) = 7.16, *p* < .0001), showing higher performance (*mean wACC* = .195) with multiple rule sets than without (*mean wACC* = .110). Moreover, performance was also significantly better (*t*(34) = 4.45, *p* < .0001) in the Stepwise environment, for models with multiple rule sets (*mean wACC* = .195) than without this feature (*mean wACC* = .110).

The strongest main effect (in terms of Cohen’s d effect size; see Table 2) however was observed for the hierarchical learning feature (χ^2^(1, *N* = 407) = 136.61, *p* < .0001), showing that also for this feature the models performed significantly better overall when they used hierarchical learning (*mean wACC* = .232) than when they did not (*mean wACC* = .134). As illustrated in Fig. 8E, and supported by a significant interaction effect (χ^2^(2, *N* = 407) = 117.783, *p* < .0001), the performance benefit strongly differed across environments. Indeed, while there was no significant performance benefit in the Stable (*t*(165) = .291, *p* = .771) or the Stepwise environment (*t*(34) = .716, *p* = .479), the models that implemented hierarchical learning (*mean wACC* = .292) clearly outperformed models that did not use hierarchical learning (*mean wACC* = .104) in the Reversal environment (*t*(205) = 11.761, *p* < .0001).

To also investigate the added value of each individual hierarchical extension in terms of model fit, the linear mixed-effects regression was also performed with wAIC as dependent variable. Also for this analysis, the statistics are summarized in Table 3 but discussed more in detail below. In the Supplementary materials we also present results for analyses with wLL and wBIC as dependent variable but for simplicity, we restricted analyses to one measure of fit in the main text.

The right column of Fig. 8 (panels B, D and F) shows the interactions between environment and hierarchical extensions in terms of wAIC. All three interactions reached statistical significance. Although the models without adaptive learning rate (mean wAIC = .234) fitted better (χ^2^(1, *N* = 407) = 60.553, *p* < .0001) on average than the models with adaptive learning rate (mean wAIC = .099) across all three environments, the interaction between adaptive learning rate and environment (Fig. 8B) did reach significance (χ^2^(2, *N* = 407) = 12.706, *p* = .012). A more in-depth exploration via pairwise t-tests revealed that the difference in wAIC between models that did or did not include an adaptive learning rate was significant in all three environments. However, as can be observed in Fig. 8B, the difference was less strong (see also Table 3 to compare effect sizes) in the Reversal environment (*t*(205) = -2.585, *p* = .0104) compared to the Stable (*t*(165) = -7.052, *p* < .001) and the Stepwise environments (*t*(34) = -3.690, p = .0008). In contrast to the model performance results, there was thus a disadvantage for the adaptive learning rate for fitting behavior in the Reversal environment. Nevertheless, the interaction between adaptive learning rate and Reversal environment followed the same pattern in behavior and performance.

Also for multiple rule sets, there was a significant main effect (χ^2^(1, *N* = 407) = 14.587, *p* =.0001), showing a better fit overall for models that use multiple rule sets (*mean wAIC* = .170) than for models that do not (*mean wAIC* = .161). The interaction between multiple rule sets and environment (χ^2^(2, *N* = 407) = .4860, *p* = .784) did not reach significance. However, this absence of significance for this interaction can again be explained by the fact that the linear mixed model also includes the hierarchical learning feature which is always combined with multiple rule sets and hence explains largely the same variance. Indeed, pairwise t-tests aimed to explore the interaction between environment and multiple rule sets indicated that the benefit in model fit for models that use multiple rule sets did not reach significance in every environment. As was also the case for model performance, the benefit of multiple rule sets is mainly driven by the Reversal environment. While the effect was not significant in the Stable (t(165) = .690, p = .491) or the Stepwise environment (t(34) = 1.385, p= .175), it did reach significance in the Reversal environment (t(205) = 7.551, p < .0001).

The main effect of hierarchical learning also reached significance (χ^2^(1, *N* = 407) = 136.61, *p* <.0001), demonstrating an improved fit when the model included hierarchical learning (mean wAIC = .210) compared to models that did not include this feature (mean wAIC = .145). However, there was also again a significant interaction effect of Hierarchical learning with the environment (χ^2^(2, *N* = 407) = 117.783, *p* <.0001). As shown in Fig. 7E, the benefit of Hierarchical learning was only observed in the Reversal environment (t(205) = 4.313, p < .0001). In the Stable environment there was a significant effect in the other direction (t(165) = -2.235, p = .027), showing better fit for the models that did not use hierarchical learning (mean wAIC = .187) compared to models that included this feature (mean wAIC = .157). In the Stepwise environment there was no significant effect (t(34)

= .611, p = .545). Hence, again a very similar pattern was found for model fit as for model performance.

## Discussion

Although flat learning via the RW rule has proven very valuable to describe human (and animal) learning (Glimcher, 2011; Rescorla & Wagner, 1972; Steinberg et al., 2013), a wide range of previous work has argued that flat learning is insufficient to capture human learning in complex environments (Bai et al., 2014; Bouchacourt et al., 2022; Liu et al., 2022; McGuire et al., 2014; Verbeke & Verguts, 2019). Therefore, several hierarchical extensions to the flat learning approach have been proposed in several different environments and datasets (Bai et al., 2014; Behrens et al., 2007; Foucault & Meyniel, 2021; Kruschke, 2008; Mathys et al., 2011; Silvetti et al., 2011; Verbeke, Ergo, De Loof, & Verguts, 2021). Crucially, an extensive and systematic evaluation of these hierarchical extensions over multiple reinforcement learning environments was lacking. Hence, it remained unclear what hierarchical extensions are needed to model human cognition and how this relates to variations in the environment. To remedy this, the current study used a meta-analytic, nested modelling approach to evaluate six models of different complexity, on three environments of different complexity. Model simulations were performed to illustrate which model performed best in each environment. Then, models were fitted to the behavioral data of ten different datasets (across the three environments). Results demonstrated that human data fitted best with the model that also performed best in that environment.

Moreover, which model fitted or performed best, varied over environments. In the Stable environments none of the hierarchical model extensions improved either the performance or the fit. For Reversal environments there was strong evidence for using multiple rule sets and hierarchical learning. More specifically, performance and fit were best for the models that combined the multiple rule sets with hierarchical learning (i.e., the Sets_learning and Full model). In the Stepwise environment, evidence was more mixed. While performance analyses revealed a benefit for using multiple rule sets, this feature did not significantly increase the fit. Nevertheless, the pattern was similar between performance and fit. Moreover, measures of fit that did not include a penalty for the number of parameters, demonstrate a significant improvement of fit by using multiple rule sets in the Stepwise environment (see Supplementary materials). Thus, current work demonstrates that humans can adaptively switch between (model) architectures depending on the environment.

One possible explanation for these results (O’Doherty et al., 2021), is that the human brain functions as a Mixture of Experts model. Here, it is suggested that the brain has different expert systems available, from which a higher-level system decides which expert receives control over behavior for the current task. Indeed, previous work has suggested that human prefrontal cortex functions as a gating network to guide behavior (Flesch et al., 2023; Miller, 2000; O’Doherty et al., 2021; Tsuda et al., 2020). Nevertheless, it is important to note that the experts in classic mixture of expert networks typically contain information on how to perform (in contrast to how to learn) a specific task. In this sense, the experts as traditionally conceived, are more closely related to the different rule sets in our hierarchical models. Critically, previous work did propose algorithms that learn how to learn (Dayan & Hinton, 1992; Duan et al., 2016; Wang et al., 2018). Also here, a hierarchical organization is proposed in which the higher-level system (again human prefrontal cortex) instructs the lower-level system on how to learn. Hence, one could consider a three-level hierarchy, with the rule sets at level 1, the rule set selection model at level 2, and the model-selecting system at level 3.

The current study considered three hierarchical extensions to the Flat model. A first extension is to add a higher-order process or parameter that adaptively manipulates lower-level parameters. Such meta-learning of lower-level parameter values has received considerable interest in models of human cognition (Silvetti et al., 2017; Wang, 2021; Wang et al., 2018) and has been linked to human medial prefrontal cortex (Khamassi et al., 2013; Wang et al., 2018), and more specifically dorsal anterior cingulate cortex (Behrens et al., 2007; Holroyd & Verguts, 2021; Silvetti et al., 2017). As mentioned in the Introduction, several different implementations of adaptive learning rates exist (Bai et al., 2014; Behrens et al., 2007; McGuire et al., 2014). While we considered the approach of (Bai et al., 2014) appropriate for the current models and tasks, future work should further investigate how different implementations of adaptive learning rate allow to better fit human data.

In terms of performance, we observed that an adaptive learning rate was beneficial (improved performance) in the Reversal environment. In terms of model fit, there was no strong support for adaptive learning rates. Nevertheless, we show in the Supplementary Materials that different learning rates are used for different datasets and environments. Additionally, we discussed before that different models are selected by adjusting the hierarchical parameters. Since weighted model fit to the human data varied over environments, there is evidence that the hierarchically higher parameters are adaptive as well. Thus, current study does not argue that humans do not adapt learning rates or other parameter values. However, rather than an online adaptation to random reversals in contingencies as we investigated in the current task, a clearer distinction (e.g., via contextual cues) might be needed between periods that require different learning rates (Simoens et al., 2023; L. Q. Yu et al., 2021). Future work might further explore when and how humans adapt lower-level parameters.

A second extension that we consider is that agents may maintain multiple rule sets. Here, the agent learns different rule sets on the lower level and on a hierarchically higher level learns/decides when to switch between rule sets. Previous work has linked this process of switching between rule sets to midfrontal theta (4-8Hz) power (Verbeke, Ergo, De Loof, Verguts, et al., 2021). In general, multiple rule sets have received increasing interest in recent years, both to model human cognition (Botvinick et al., 2009; Collins & Frank, 2013; Eckstein & Collins, 2020; Holroyd & McClure, 2015; Holroyd & Verguts, 2021; Verbeke & Verguts, 2019) and to advance artificial intelligence (Çatal et al., 2021; Chung et al., 2017; Dietterich, 2000). Indeed, keeping multiple rule sets has several computational advantages. As demonstrated in previous work (Verbeke & Verguts, 2019), storing multiple rule sets allows to avoid catastrophic forgetting because a change in stimulus-action-reward contingencies does not require to overwrite old information but to just switch to another rule set. Furthermore, learning is much faster if it is possible to select from previously learned rule sets (Botvinick et al., 2009; Dietterich, 2000). Hence, agents who store multiple rule sets are also better at generalizing old knowledge to novel tasks or contexts (Collins & Frank, 2013; Eckstein & Collins, 2020).

A third extension considered learning at the hierarchically higher level. Here, learning at the hierarchically higher level allowed to adaptively change the weight of negative feedback in the switch accumulation process. Specifically, without hierarchical learning, the model would always switch after a specific number of trials with negative feedback (e.g., after 2 consecutive trials with negative feedback). This is appropriate if variability in the feedback is consistent over time and known to the agent. However, in the current datasets, subjects were not instructed about the exact feedback reliability and hence needed to learn about the variability of the feedback. Furthermore, in the Stepwise environments, P(*Rew*|*aoptimal*) would also change during the time course of the task. Nevertheless, this did not result in a benefit for hierarchical learning for performance or fit in this Stepwise environment.

Like the adaptive learning rate, also learning at the hierarchically higher level can have multiple manifestations. For instance, instead of learning when to switch rule sets, one can also learn which rule set to select given a current task context (Botvinick et al., 2009; Holroyd & McClure, 2015). This allows to select the appropriate rule set in each context. One popular framework (Collins & Frank, 2013; Gershman & Niv, 2012; Niv, 2019) extends this approach and suggests that on top of grouping stimulus-action associations in rule sets, contexts can also be grouped in latent states. As a result, contexts are mapped to latent states which represent lower-level rule sets (one rule set for each latent state). This allows for fast generalization since lower-level knowledge (stored in rule sets) can immediately be transferred to all contexts that map to the same latent state.

Hence, previous work has proposed three levels of abstractions from rule sets at the lowest level to contexts and to latent states at the highest level. It has been suggested that these levels of abstraction are organized along the rostral-caudal axis of human prefrontal cortex where latent states are represented at the most rostral parts of prefrontal cortex (Badre et al., 2010; Badre & Nee, 2018; Boorman et al., 2009; Koechlin et al., 2003; Nakahara et al., 2002).

Strikingly, both the adaptive learning rate and hierarchical learning extension seem especially adaptive in noisy environments with varying levels of uncertainty. On the lower level of the rule sets, adaptive learning rates could be employed to balance behavioral adaptation in response to two types of uncertainty; uncertainty about feedback validity when P(*Rew*|*a_optimal_*) < 100% but also uncertainty about the identity of *a_optimal_* in environments where this can change (Reversal and Stepwise). Previous work has linked these two types of uncertainty to the neuromodulators norepinephrine and acetylcholine (Aston-Jones & Cohen, 2005; A. J. Yu & Dayan, 2005). On the higher level, hierarchical learning could be employed to adapt rule switching to the uncertainty about feedback validity. When this validity is low, the agent should be more conservative to switch between rule sets than when this validity is high.

To evaluate the three hierarchical extensions, current work adopted a nested modelling approach (Grainger & Jacobs, 1996; Perry et al., 2007; Vidal & Durfee, 1998). Specifically, six models were constructed by considering different combinations of the three hierarchical extensions (see Fig. 1A). Importantly, such a nested approach allows to investigate the added value of each individual extension but also of their interactions. Nevertheless, it is important to note that for each hierarchical extension, different computational implementations exist in literature. Hence, future work should investigate how such different implementations could lead to different results. Moreover, several other extensions could be made to the current models. For instance, as suggested by the GRU network (see Fig. 6B), the models could benefit from having more than two rule sets. Potentially each rule set could in that case also benefit from having a separate learning rate or temperature parameter. A significant improvement could be made by extending the model in a way that it can itself select and tune the learning parameters and algorithm that is appropriate in the current environment (Duan et al., 2016; Wang et al., 2018). One option is to use Bayesian inference to select the best learning algorithm (Tavoni et al., 2022).

All six models were evaluated on three environments. In Stable environments, the stimulus-action-reward contingencies did not change. Hence, the agent must learn one single rule set. For this Stable environment, the Flat model was optimal. As described in the Introduction, high learning rates would be suboptimal in noisy environments since it would lead the agent to “chase the noise”. In the Supplementary Materials, we show optimal parameter values and estimated parameters for each dataset. Here, it is indeed observed that for more noisy datasets (lower P(*Rew*|*a_optimal_*)), lower learning rates lead to better performance and fit behavioral data better. Furthermore, in the noisiest (Goris/Stable) dataset, we also observed a better fit for the Sets model which could indicate that instead of integrating trial-by-trial feedback into one set of stimulus-action mappings, subjects create two alternative hypotheses (in rule sets) and alternate between those two options. Together, this suggests that Flat learning models experience significant problems if they are faced with noisy environments that require strong flexibility.

To test this, a second environment considered reversal learning tasks. Results demonstrated that the Flat model is not suited for Reversal environments. Here, the Sets_learning and Full models performed best, and also fitted best to human behavioral data. As supported by the linear mixed models, there was thus a strong advantage for models that use both multiple rule sets and hierarchical learning. Importantly, models with only multiple rule sets (Sets and Sets_ALR model) did not perform better than the Flat model. This suggests that models were better to decide when to switch between rule sets when they could also learn at the hierarchically higher level. As described before, learning on the hierarchically higher level is beneficial when reliability of feedback changes over time. At first sight, this seems particularly relevant for the Stepwise environment. However, in the current tasks where stimulus-action-reward contingencies needed to be learned from scratch, the reliability of feedback is actually dynamic in every environment. Specifically, feedback is always more variable when the appropriate lower-level rule set is not learned yet compared to when learning on the lower level has converged. Hence, while receiving negative feedback during the initial learning of a rule set probably just signals that the agent should learn this rule set better, negative feedback after learning of the rule set could signal an actual change in stimulus-action-reward contingencies. Therefore, it is better to have learning on the hierarchically higher level in the Reversal environment as well.

Notably, the Goris/Volatile dataset seemed to be the only dataset in which a better fit was observed for models that used multiple rule sets but no hierarchical learning. However, Fig. 2A shows that accuracy was also significantly lower for this dataset. One causal factor for this lower accuracy could be that for the Goris/Volatile dataset, reversals occurred on average after 18 trials with fastest reversals occurring after only 15 trials in this dataset. In other datasets there was considerably more time between two reversals (average reversals after at least 30 trials for tasks with P(*Rew* |*a_optimal_*) < 100%; see also Table 1). Hence, the fast succession of reversals in the Goris/Volatile dataset may have caused significant problems for subjects (as evidenced by a low accuracy) and caused subjects to drop the hierarchical learning extension.

The Full model contained all three hierarchical extensions and hence added an adaptive learning rate on top of the multiple rule sets and hierarchical learning. Although evidence for the ALR model was consistently low across datasets, there seemed to be a benefit for the adaptive learning rate in the Reversal environment when it is combined with multiple rule sets and hierarchical learning. One possible explanation could be that the ALR model needs to adapt the learning rate in two directions. More specifically, it needs to decrease the learning rate in stable periods and increase the learning rate when reversals occur. For models with multiple rule sets, this increase of the learning rate on reversals is not necessary since it can just switch between rule sets on reversals. Hence, for the Full model, the learning rate should be high only when the model learns a certain rule set for the first time. Once it has learned the rule, it needs stability (robustness against noise) and hence the learning rate can decrease. If later a reversal occurs, the Full model does not have to increase the learning rate again but can just switch to another rule set.

Whereas in the Reversal environment the identity of the optimal action could regularly change, P(*Rew*|*a_optimal_*) was stable across the entire task. Hence, we considered a third environment in which P(*Rew*|*aoptimal*) could change as well. We referred to this environment as the Stepwise environment. Model simulations indicated that also here model performance would benefit from using multiple rule sets. Although evidence in the empirical datasets was more mixed, model fit (Fig. 7) and mixed model regression (Fig. 8D) shows a small advantage for models with multiple rule sets relative to those without multiple rule sets. Although regression results illustrated that this was not significant, the effect size of d = .238 (see Table 3) suggest that the absence of effect could be due to a lower power with only 35 subjects in this environment.

Furthermore, for both datasets in the Stepwise environment, there were periods in which P(*Rew*|*a_optimal_*) = 50%. This is highly ambiguous since there is in fact no optimal action, and it is unclear for the model which rule set should receive control over behavior. As suggested before, one option for modeling would be to include additional rule sets. However, this would add another layer of complication, as it would mean that the model should either randomly switch between the rule sets, or instead implement a decision mechanism allowing to infer to which rule set it should switch to. Hence, future work should investigate whether our models are generalizable to Stepwise environments or additional extensions should be made. Alternatively, future work might consider evaluating model evidence in Stepwise environments without this P(*Rew*|*a_optimal_*) = 50% condition.

In sum, while flat learning strategies are optimal for Stable learning environments, current work found support for hierarchical extensions in the Reversal environment. While Flat models need to overwrite rule sets at each rule switch, hierarchically organized models can store multiple (stable) rule sets and on a hierarchical higher level, and flexibly decide when to switch between rule sets. This provides an important computational advantage in Reversal environments; an advantage that humans also seem to exploit. Moreover, we found an additional improvement in fit and performance when the model was further extended with learning on the hierarchically higher level and an adaptive learning rate on the lower level. The added value of these features can be explained by the fact that rule sets should still be learned at initial stages of the task. This means that the model should learn faster and be more conservative for rule switching during initial rule learning than once rule sets are learned. While model simulations predicted that multiple rule sets would also be beneficial in Stepwise environments, there was no significant evidence for this extension in the empirical datasets.

Critically, current study was quite restrictive in its selection of datasets. Here, two factors were investigated; the reinforcement environment (Stable, Reversal or Stepwise) and the noise level (defined as P(*Rew* |*a_optimal_*)). We aimed to keep all other factors as consistent as possible across datasets. Hence, an important avenue for future research is to investigate how other task features influence the performance and behavioral fit of our models. For instance, while all datasets here were 2-armed bandits, increasing the number of arms would also require more than 2 rule sets. Additionally, both arms were always anti-correlated, meaning that a change in reward probability for one action always meant that the reward probability for the other action also changed in a perfectly anti-correlated manner. In these situations, it is quite beneficial to store stimulus-action mappings in rule sets because if one mapping changes, all change in a predictable manner. However, if the reward probability for each action changes in an uncorrelated manner, the Flat model might be better again. Additionally, one could investigate continuous reinforcement dynamics on top of the discrete Reversal or Stepwise changes that were made in current datasets.

We observed that the GRU network performed the tasks very well; but in a sense it did too well, as it outperformed both the humans and the cognitive models. Critically, compared to neural networks, human agents have much more limited computational resources that they can assign to a novel task (Lieder & Griffiths, 2020). Therefore, they apply heuristics which are better described by the cognitive models. Notably, as an alternative to our “bottom up” model construction starting from the flat model, one could take a “top down” approach and start from the GRU (or another recurrent network) and systematically add restrictions to it in order to investigate how it can correspond to human data. This is an avenue for future research (but see also Miller et al., 2023).

Meta-analysis has proven to be a valuable tool to address the replication crisis (Carter et al., 2019; Maxwell et al., 2015; Open Science Collaboration, 2012; Shrout & Rodgers, 2018). As we also observed across datasets in the Reversal and Stepwise environments, multiple single studies can often have varying, or even contradicting, results. Additionally, previous work has raised the issue that computational models tend to generalize poorly across different tasks (Eckstein et al., 2021, 2022). Nevertheless, over many datasets, a clear pattern can arise which allows to make more reliable conclusions. Importantly, as in all meta-analyses, one should consider the publication bias (Thornton & Lee, 2000). Because of this publication bias, possibly relevant datasets were overlooked if they were not published. However, a strength of the current study is that the datasets that were used here were not previously used to test the same models and hypotheses as in the current analyses. Hence, the publication bias that might be present in the selection of datasets is not specific to our hypotheses.

As is demonstrated in current work, and previously also in the context of working memory (van den Berg et al., 2012), meta-analyses can be useful for model selection. A good model of human cognition must fit human data better than an alternative model. However, most work typically considers such a model comparison for one (or a few) dataset(s) and one cognitive problem (task). This is suboptimal because, as we have demonstrated, model evidence can differ across learning problems and even across multiple datasets investigating the same learning problem. Therefore, we carried out model comparison over multiple datasets and learning environments. This allows to gain a deeper insight in which computational features are relevant for fitting human behavior and how these features relate to different environments. Specifically, we demonstrated that in Stable learning environments, Flat learning via the RW rule is sufficient to fit human behavior. However, for more complex environments in which stimulus-action-reward contingencies can reverse or change in a stepwise manner, we find evidence for more complex models that implement hierarchical extensions of the Flat learning approach. Particularly a hierarchical model architecture which stores multiple rule sets at the lower level and decides at a higher level which rule set should receive control over behavior, is beneficial in these environments. We believe there is significant potential in extending our meta-analytic approach with different models and different environments.

## Acknowledgements

We thank Pieter Huycke, Liyu Xia, Judith Goris, Meng Liu, Qi Chen, Dahlia Mukherjee, Michael X. Cohen and Thomas P. Hein for sharing their datasets either via open science framework or via direct communication with us. Additionally, we thank Juliet Davidow to share her stimuli with us to use for the collection of the online additional dataset. We also thank Senne Braem for helping us interpret some of the datasets. PV and TV were both supported by the Research Foundation Flanders (FWO) / Fonds Nationale de la Reserche Scientifique EOS grant G0F3818N. Additionally, PV was supported by grant 1102519N from Research Foundation Flanders. We thank Irene Cogliati-Dezza and Thomas Colin for their comments on an earlier version of the paper. Furthermore, we thank three anonymous reviewers for their constructive feedback.

## Code and data availability statement

All code as well as the behavioral data that we collected is available on our GitHub repository: https://github.com/CogComNeuroSci/PieterV_public/tree/master/Model_Meta. For data from previous publications that we included in our analyses, we refer to the original manuscripts for links to repositories or contact information of the corresponding authors.

## Notes

### Competing Interest Statement

The authors have declared no competing interest.

### Summary of Updates

Four major changes are made to the manuscript. First, we now optimized (in the sense of finding parameters that yield the best performance) all models on each dataset separately. This made the model performance results more comparable to the model fit results. This also allowed us to apply the same linear mixed model to model performance and model fit. Second, we implemented a recurrent neural network (GRU) as normative benchmark for optimality of performance. This GRU network informed us that humans are not optimal; but neither are the cognitive models, which are in fact closer to the humans than they are to the GRU. Third, we provide more insights in the datasets, models and statistical results. A table now provides more details for each dataset; model and parameter recovery results are moved to the main manuscript; learning curves and model dynamics are illustrated in novel figures; and tables are provided for statistical results. Fourth, we adapted the discussion to provide a more nuanced framing of our work in previous literature.

