## Supplementary materials for "Humans adaptively select different computational strategies in different learning environments"

#### Regression on other measures of fit

In the main text, we reported results of a linear mixed model with wAIC for each subject in each dataset as dependent variable. Here, environment (Stable, Reversal or Stepwise), adaptive learning rate (yes or no), multiple rule sets (yes or no) and hierarchical learning (yes or no) were included as fixed independent variables. Dataset was included in the analyses as a random factor (intercept). We chose wAIC because it provides a balanced penalty for the number of parameters in the model. However, here we also report results for this linear mixed model when no penalty is applied (wLL) and when a more severe penalty for the number of parameters is applied (wBIC). Fig. S1 demonstrates the interaction effect between computational features and the environment and Table S1 reports statistical results. We observe a very robust advantage of combining multiple rule sets with hierarchical learning irrespective of the penalty for the number of parameters. Additionally, the results of wLL show a stronger similarity to the results of wACC shown in Fig. 8 A, C and E in the main text. Note that this wACC measure also does not include a penalty for the number of parameters.

**Table S1. Statistics.**

**wLL**

| | *Linear mixed model* | | *Pairwise t-tests* | | |
|---|---|---|---|---|---|
| | Main effect | Interaction with environment | Stable environment | Reversal environment | Stepwise environment |
| *Adaptive learning rate* | No > Yes<br>*p* = .484<br>*d* = .029 | *p* = .003 | No > Yes<br>*p* = .056<br>*d* = .150 | Yes > No<br>*p* = .186<br>*d* = .093 | No > Yes<br>*p* = .057<br>*d* = .338 |
| *Multiple rule sets* | Yes > No<br>*p* < .0001<br>d = .837 | *p* = .451 | Yes > No<br>*p* < .0001<br>*d* = .835 | Yes > No<br>*p* < .0001<br>*d* = .857 | Yes > No<br>*p* < .0001<br>*d* = 1.187 |
| *Hierarchical learning* | Yes > No<br>*p* < .0001<br>*d* = .596 | *p* < .0001 | Yes > No<br>*p* < .0001<br>*d* = .370 | Yes > No<br>*p* < .0001<br>*d* = .795 | Yes > No<br>*p* = .006<br>*d* = .496 |

**wBIC**

| | *Linear mixed model* | | *Pairwise t-tests* | | |
|---|---|---|---|---|---|
| | Main effect | Interaction effect | Stable environment | Reversal environment | Stepwise environment |
| *Adaptive learning rate* | No > Yes<br>*p* < .0001<br>*d* = .895 | *p* < .0001 | No > Yes<br>*p* < .0001<br>*d* = 1.463 | No > Yes<br>*p* < .0001<br>*d* = .577 | No > Yes<br>*p* < .0001<br>*d* = 2.071 |
| *Multiple rule sets* | No > Yes<br>*p* < .0001<br>*d* = .237 | *p* < .0001 | No > Yes<br>*p* < .0001<br>*d* = .687 | Yes > No<br>*p* = .022<br>*d* = .161 | No > Yes<br>*p* < .0001<br>*d* = .896 |
| *Hierarchical learning* | No > Yes<br>*p* = .579<br>*d* = .138 | *p* <.001 | No > Yes<br>*p* < .0001<br>*d* = .940 | Yes > No<br>*p* = .003<br>*d* = .210 | No > Yes<br>*p* = .020<br>*d* = .418 |

*Notes.* Two types of statistical tests were performed. The first statistical test was the linear mixed model (left part). The second type of statistical tests (right part) were pairwise t-tests. These t-tests provide a better insight into the interaction effect from the linear mixed model. Both types of statistical tests were performed with either wLL (upper part) or wBIC (lower part) as independent variable. For the main effect of the linear mixed model as well as for the pairwise t-tests a direction of the difference is reported. This direction indicates whether the model with a hierarchical extension (in the columns) scores better than the model without that hierarchical extension (Yes > No) or the other way around (No > Yes). A p-value indicates the significance of this difference and the Cohen's d effect size provides an indication of how strong this effect is. For the interaction effect, we only provide a significance (p-) level.

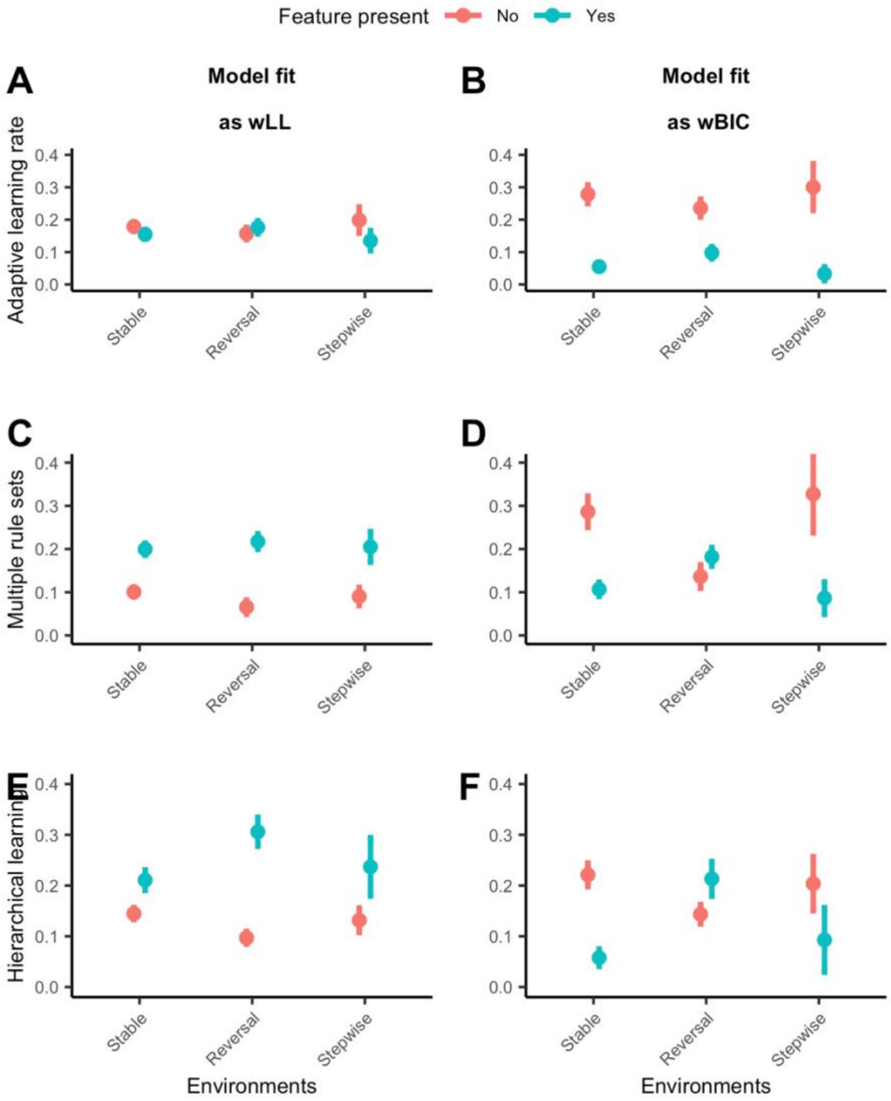


**Figure S1. Additional regression results for hierarchical model extensions in each environment.** Each datapoint reflects the mean (wLL or wBIC) over all models with one particular feature (e.g., the ALR, Sets_ALR and Full model for the adaptive learning feature) in a particular environment. Column 1(A, C, E) illustrates the influence of each extension on model fit, measured as wLL (on y-axis), for the behavioral datasets in each environment. . Column 2(B, D, F) illustrates influence of each extension on model fit, measured as wBIC (on y-axis), for the behavioral datasets in each environment.

### Model dynamics

Fig. S2 and S3 provide a more detailed insight in the model dynamics. For illustration purposes, we selected the Mukherjee dataset which consisted of 90 trials, evenly divided in three blocks of 30 trials following an ABA rule sequence. Although the exact rule was counterbalanced across subjects, we coded it here such that a left response was optimal on the first and last 30 trials (rule A) and the right response was optimal for the second 30 trials (rule B). In this dataset, reward feedback was very noisy: P($Rew|a_{optimal}$) = 70 %.

Fig. S2 demonstrates the fit of each model (rows) for three subjects (columns). Here, subjects were selected as the percentile 0 (A, D, G, J, M & P) , 50 (B, E, H, K, N, Q) and 100 (C, F, I, L, O, R) in terms of overall task accuracy. We show subject responses (black crosses) and likelihood of a right response as predicted by the model (red line).

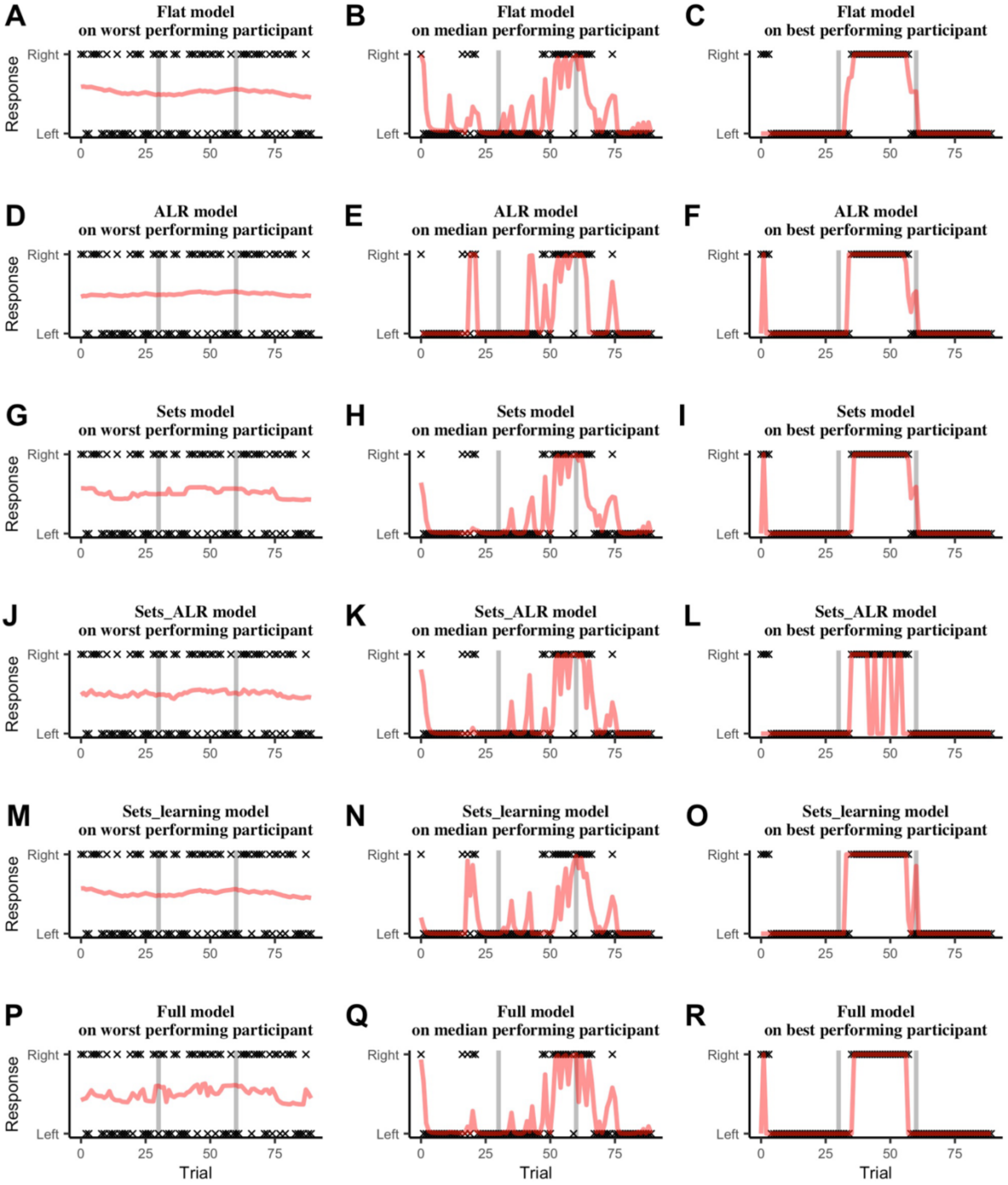


**Figure S2. Fit dynamics of each model on three subjects of the Mukherjee dataset.** Subjects were selected as the percentile 0 (A, D, G, J, M & P) , 50 (B, E, H, K, N, Q) and 100 (C, F, I, L, O, R) in terms of overall task accuracy. Black crosses indicate subject responses, and the red line illustrates the likelihood of a right response as predicted by the model.

Fig. S3 demonstrates the performance for each model (rows) for three different simulations (columns). Here, simulations were performed on all subject designs. Hence also performance of the models varied over the designs. Again, we selected the 0, 50 and 100 percentiles in terms of overall task accuracy. Here, the blue line represents the correct response, and the red crosses represent the likelihood that the model gave the right response as given by the softmax activation function given by Equation (2) in the main text.

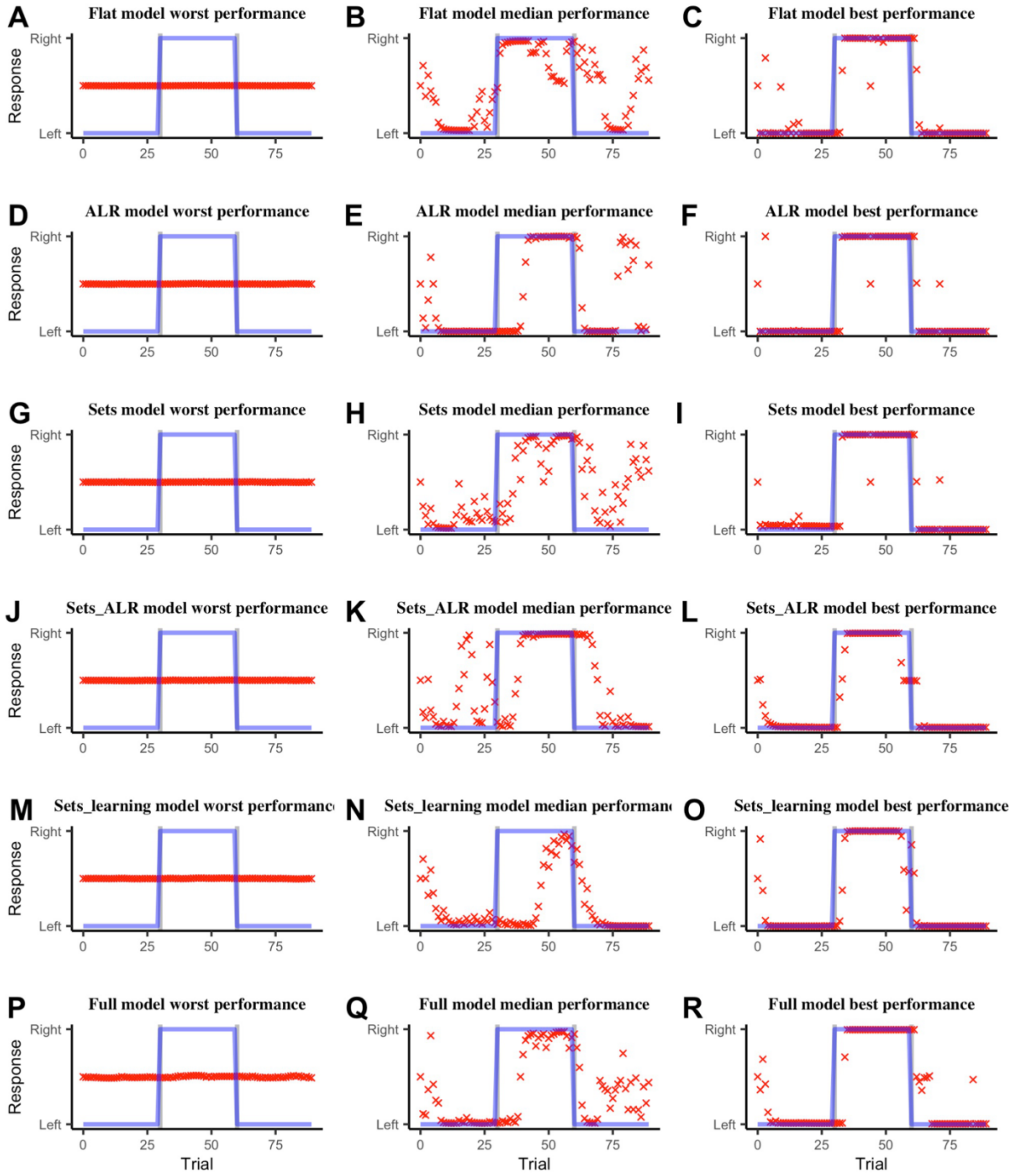


**Figure S3. Performance dynamics of each model on three simulations of the Mukherjee dataset.** Simulations were selected as the percentile 0 (A, D, G, J, M & P) , 50 (B, E, H, K, N, Q) and 100 (C, F, I, L, O, R) in terms of overall task accuracy. The blue line indicates the most optimal response and the red crosses illustrates the likelihood that the model gave the right response as given by the softmax activation function given by Equation (2) in the main text.

**Parameter results**

We present parameter values for all models in all environments and/or datasets. Fig. S3 provides parameter values as optimized for performance (in terms of accumulated reward) in each environment. Fig. S4 provides (scaled) parameter values as optimized in terms of model fit to empirical data from the datasets. Values are scaled to allow for easier comparison.

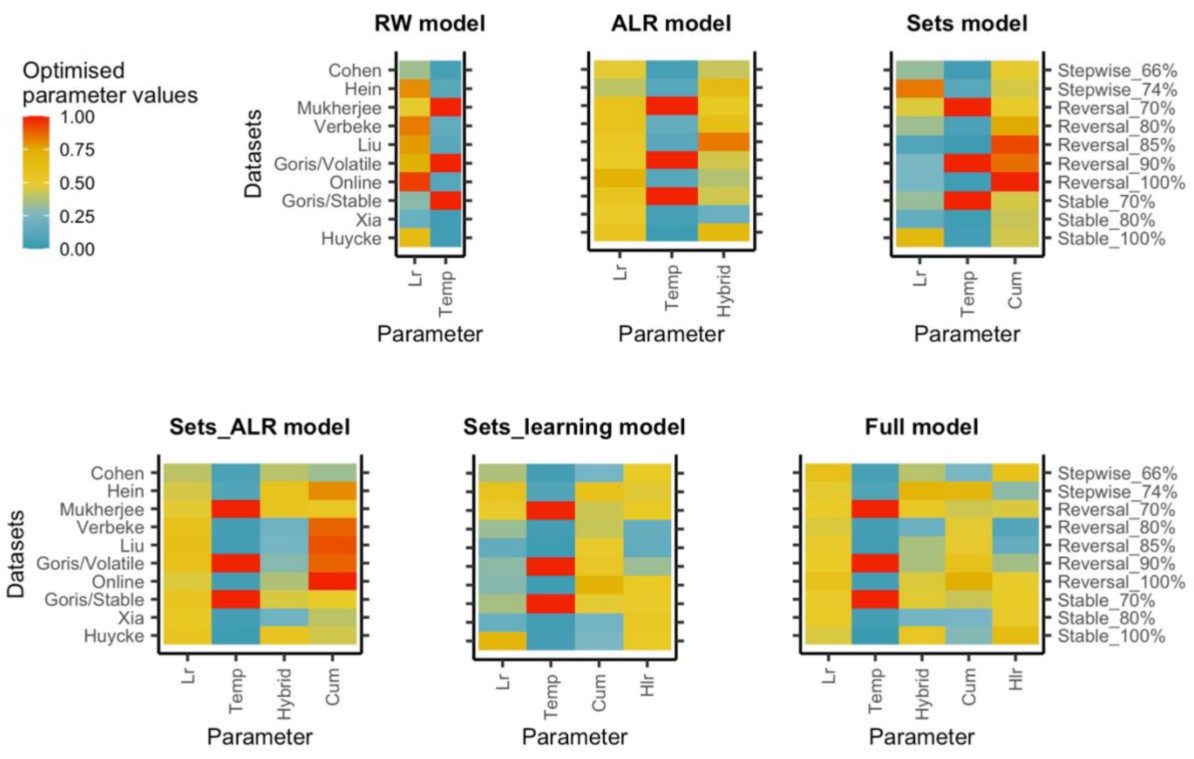


**Figure S4. Optimized parameter values for each model in each environment.** Parameters were optimized based on accumulated reward across all trials in each dataset.

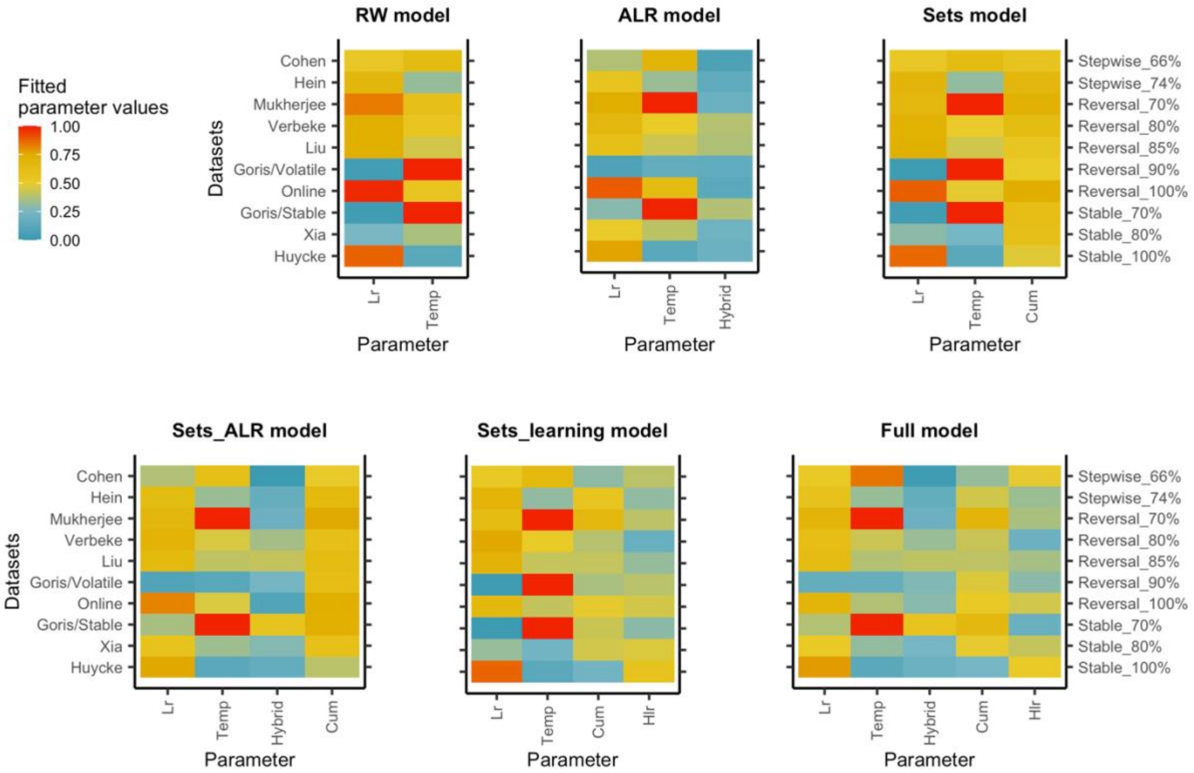


**Figure S5. Estimated parameter values for each model in each dataset**. Parameters were estimated by minimizing the negative log likelihood of response via differential evolution.
